# Neuromodulators generate multiple context-relevant behaviors in a recurrent neural network by shifting activity flows in hyperchannels

**DOI:** 10.1101/2021.05.31.446462

**Authors:** Ben Tsuda, Stefan C. Pate, Kay M. Tye, Hava T. Siegelmann, Terrence J. Sejnowski

**Affiliations:** Computational Neurobiology Laboratory, The Salk Institute for Biological Studies, La Jolla, CA 92037, USA; Neurosciences Graduate Program, University of California San Diego, La Jolla, CA 92093, USA; Medical Scientist Training Program, University of California San Diego, La Jolla, CA 92093, USA; Systems Neuroscience Laboratory, The Salk Institute for Biological Studies, La Jolla, CA 92037, USA; Biologically Inspired Neural & Dynamical Systems Laboratory, School of Computer Science, University of Massachusetts Amherst, Amherst, MA, 01003; Institute for Neural Computation, University of California San Diego, La Jolla, CA 92093, USA; Division of Biological Sciences, University of California San Diego, La Jolla, CA 92093, USA

## Abstract

Neuromodulators are critical controllers of neural states, with dysfunctions linked to various neuropsychiatric disorders. Although many biological aspects of neuromodulation have been studied, the computational principles underlying how neuromodulation of distributed neural populations controls brain states remain unclear. Compared with specific contextual inputs, neuromodulation is a single scalar signal that is broadcast broadly to many neurons. We model the modulation of synaptic weight in a recurrent neural network model and show that neuromodulators can dramatically alter the function of a network, even when highly simplified. We find that under structural constraints like those in brains, this provides a fundamental mechanism that can increase the computational capability and flexibility of a neural network. Diffuse synaptic weight modulation enables storage of multiple memories using a common set of synapses that are able to generate diverse, even diametrically opposed, behaviors. Our findings help explain how neuromodulators “unlock” specific behaviors by creating task-specific hyperchannels in the space of neural activities and motivate more flexible, compact and capable machine learning architectures.

**Significance:** Neuromodulation through the release of molecules like serotonin and dopamine provides a control mechanism that allows brains to shift into distinct behavioral modes. We use an artificial neural network model to show how the action of neuromodulatory molecules acting as a broadcast signal on synaptic connections enables flexible and smooth behavioral shifting. We find that individual networks exhibit idiosyncratic sensitivities to neuromodulation under identical training conditions, highlighting a principle underlying behavioral variability. Network sensitivity is tied to the geometry of network activity dynamics, which provides an explanation for why different types of neuromodulation (molecular vs direct current modulation) have different behavioral effects. Our work suggests experiments to test biological hypotheses and paths forward in the development of flexible artificial intelligence systems.

## Introduction

Neuromodulators are a central mechanism in the biological control of neural states that manifest as mood, arousal, motivation and other variable behavioral modes [1–8]. Other mechanisms, like exogenous contextual cues (e.g. a red light at a traffic signal), can also generate specific behaviors [9] by providing input signals that elicit specific activity patterns in a neural network. Although exogenous cue-based mechanisms can be advantageous in some circumstances (e.g. fast behavioral switching), they are impractical when massive scaling is needed. Broadcasting cue information across large brain regions would require many millions of neural connections to simply convey a constant cue signal to maintain conditioned activity [10]. Neuromodulation is an ideal system to control major changes of behavioral state, albeit at slower timescales, by modifying how neurons transduce information, including modulation of intrinsic ion channels and synaptic strengths. The impact of neuromodulation is not as specific as arrays of heterogeneous inputs, but we show here that it can flexibly select different computational outcomes in response to the same heterogeneous inputs.

A classic example of this trade-off is illustrated by sleep. Humans and many other species cycle through internal sleep stages that are controlled by a variety of neuromodulators. An exogenous cue-dependent system would require a “sleep cue” to be widely broadcast across the whole brain, constantly for hours, at great energetic expense. In contrast, neuromodulators that broadcast sparse signals widely can shift the entire brain into internal sleep states that persist for hours [3, 8].

The diffuse release of neuromodulators from a relatively small number of neurons projecting across huge regions of the brain can control a variety of “internal states” [5, 8, 11–13]. In addition to sleep, neuromodulators are responsible for controlling mood, states of alertness [8, 14] and regulating the consumption of food and water during awake states [1, 7]. Pioneering studies on the lobster and crab pyloric networks [15–17] and other systems [18, 19] have revealed how neuron-specific neuromodulation can precisely tailor central pattern generator rhythms. Yet it remains unknown how large-scale neuromodulation of vast distributed neural populations can control global network dynamics and dictate behavior as it does in large brains.

Fully understanding neuromodulation in brains is important for several reasons. First, most psychiatric disorders either stem from or are directly related to neuromodulator dysregulation, and many psychiatric drugs target neuromodulatory activity [5, 20, 21]. Second, many of the psychiatric drugs currently only partially or imprecisely target neuromodulatory processes and effect their results through mechanisms that remain unclear [4, 22, 23]. Third, effects of many psychiatric treatments are highly variable, with some patients responding strongly and others failing to respond to multiple drugs [20, 24–26]. Fourth, neuromodulation acts via multiple mechanisms (as discussed below), allowing powerful circuit control but also making it difficult to fully understand their combinatorial actions [5, 27, 28]. Fifth, given the central role of neuromodulators in control of brains, a better understanding promises to make deep learning models based on brain architectures more flexible, more compact, and more efficient.

Neuromodulators affect several processes in brains including synaptic strengths, neural excitability, plasticity, and, indirectly, downstream circuit activity [15, 28–30]. Prior research has focused on different aspects of neuromodulation [31], including Yu and Dayan [32] who modeled the role of acetylcholine and norepinephrine in Bayesian probability estimations of uncertainties; Stroud et al. and Vecoven et al. [33, 34] who considered modulation of the neural activation function; Beaulieu et al. [35] who formulated neuromodulation as a separate network that masks effector networks; Miconi et al. [36] who used modulation of synaptic plasticity to train networks; and Hasselmo et al. [37] who developed a model incorporating experimental work on multi-factor neuromodulator-specific circuit dynamics, particularly in hippocampal memory processes. Our model of synaptic weight modulation shares some similarities to previous models, particularly to the neural excitability models [33, 34]. Both lead to increased flexibility and versatility, yet they operate through independent mechanisms both biologically [28] and computationally (see Extended Data Appendix A).

We focus here on a critical aspect of neuromodulation in brains — synaptic weight modulation [15, 17, 18, 27, 28, 38] — a poorly understood and important control mechanism in brains. We consider a simplified approximation in which neuromodulators act as nearly uniform synaptic weight amplifiers or dampeners — a single multiplicative scalar — within a local region of a neural network. We show how this form of neuromodulation establishes distinct memories using a common set of synapses, generating unique dynamic activity landscapes within a structurally-conserved neural network. We demonstrate how neuromodulated circuits give rise to idiosyncratic, non-linear dose-response properties that can differ depending on the mode of neuromodulation. Using a neuromodulation-mediated behavioral paradigm in *Drosophila*, we show how this form of neuromodulation naturally handles intermediate neural states, and as such, generalizes models of discrete internal state switching [39, 40] to continuous state transitions. Although many mechanisms may influence behavioral shifts, we show that a simple multiplicative factor applied to weights already acts as a powerful network control device, allowing neuromodulators to vastly increase the capability and complexity of computation in brains and making artificial neural networks more flexible, compact and capable.

## Results

### Neuromodulation creates multiple weight regimes within shared synaptic connections

The effects of neuromodulators on synaptic weights present a mode of circuit control [28] that is poorly understood in brains — both how it is implemented at scale and the computational mechanisms by which it shifts coordinated activity to generate different behaviors. Several recent studies on cell type diversity have made clear that brains contain a complex array of neuromodulators that act with carefully coordinated spatiotemporal precision [41–45]. As a first step, we sought to assess whether a simplified form of neuromodulation — modeled as a uniform multiplicative factor acting on synaptic weights in a recurrent neural network (RNN) — could help us understand how neuromodulators control neural state. Although other modes of neural network control such as exogenous contextual cuing have been shown to successfully shift network behavior [9], uniform weight modulation represents a completely different biological and computational mechanism, which, given the complex, non-linear, and often unpredictable nature of RNNs, requires explicit assessment. Furthermore, systems in which exogenous cues are required to drive neurons [9] are unsurprisingly more sensitive to external noise (Extended Data Fig. 1), while internally controlled neuromodulation-based systems are robust to such fluctuations in external inputs [46] (Extended Data Fig. 1; although they are subject to internal sources of noise: see Extended Data Fig. 1g–j), supporting neural states like sleep as described above [3].

We designed a simple experimental paradigm which we call “Positive-Negative Output Task” (Fig. 1a; see Methods) to assess whether given identical input stimuli, uniformly shifting all the recurrent weights in a RNN could elicit a unique trained behavior from the same network. In the Positive-Negative Output Task, one of two possible stimuli (“+” or “∅”) are input to the RNN. The RNN is trained to produce either a positive, zero, or negative output for each stimuli. For example, in Fig. 1a, “Behavior 1” in black shows output from an RNN trained to produce a positive output when given the positive (+) input signal and to produce zero output when given the zero (∅) input signal. Under a different neuromodulation state, depicted in Fig. 1a as the blue RNN and “Behavior 2,” the RNN produces different outputs to the same input stimuli (e.g. + stimulus elicits a zero output and ∅ stimulus elicits a negative). This task simulates how a human or animal may behave differently to the same external cues while in different internal state. For example, a thirsty mouse cued that only water is available (+ stimulus) may run to the water (positive output) and do nothing when no water cue is given (∅ stimulus leads to zero output), but a hungry mouse may do nothing when cued that only water is available (+ stimulus leads zero output) and forage for food when no water-only cue is given (∅ stimulus leads to negative output).

We found that neuromodulation simulated as broadcast synaptic weight scaling was able to generate the distinct behaviors for the task. For example, scaling all weights by a factor of 0.5 (Fig. 1b) could generate unique output behaviors (Fig. 1c). This simple mechanism demonstrates how neuromodulators operating in brains can effectively separate synaptic memory regimes within a fixed circuit and access them through uniform scaling of weights to “unlock” specific behaviors (Extended Data Fig. 3). We found this result held over a wide range of neuromodulatory factor magnitudes (Extended Data Fig. 4a–c).

**Figure 1.**
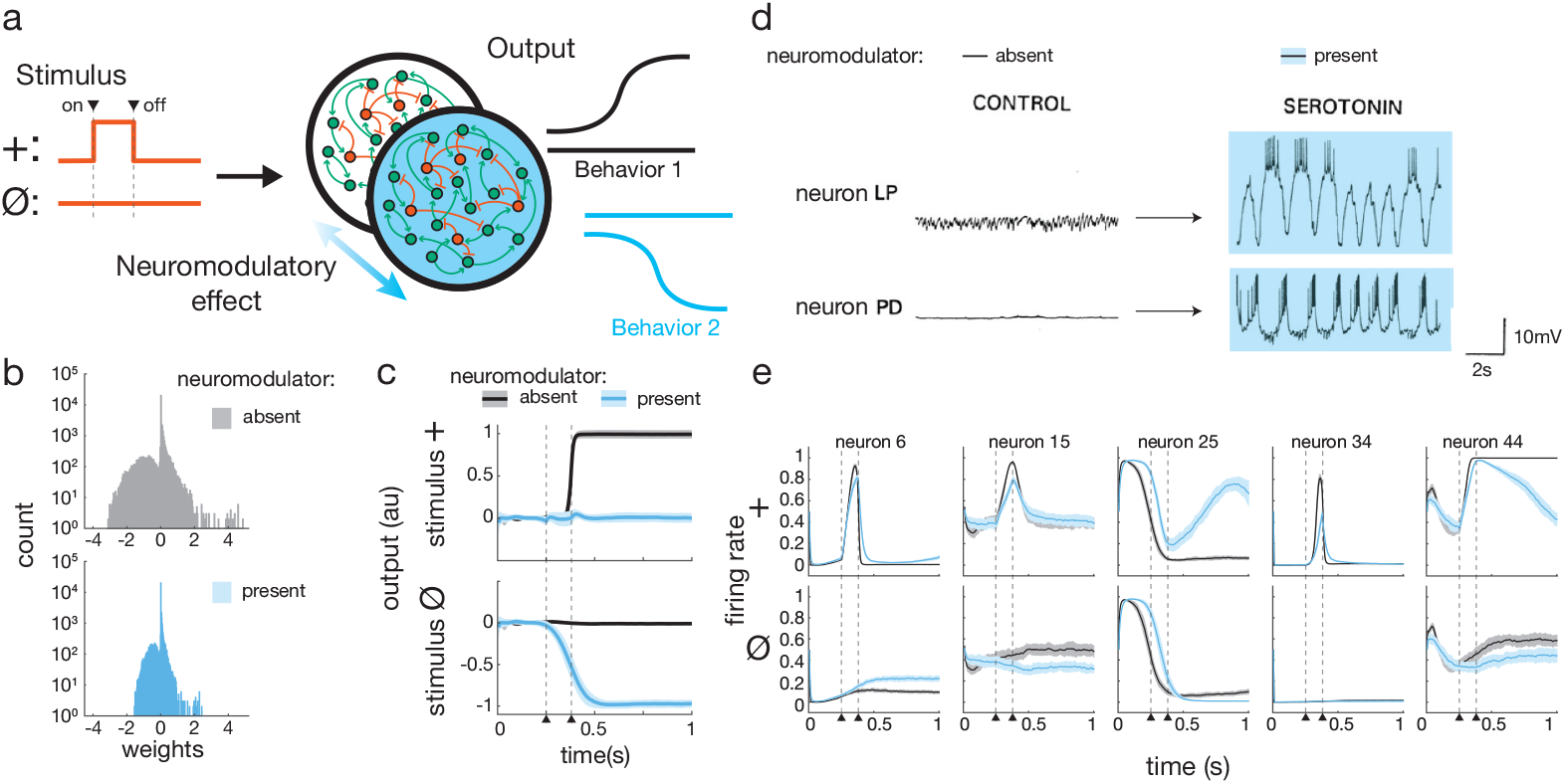
Neuromodulation weight scaling separates overlapping synaptic memory regimes. (*A*) Positive-Negative Output Task. Given a stimulus (either + or ∅), in absence of neuromodulatory effect the recurrent neural network (RNN) should produce outputs from the Behavior 1 repertoire (input + → positive output; input ∅ → zero output) and in presence of neuromodulator, from Behavior 2 (input + → zero output; input ∅ → negative output). (*B*) Neuromodulatory effect implemented in the model: all synaptic weights in the RNN are multiplied by a constant factor, here 0.5. (*C*) Mean output from 10 independently trained RNNs to + and ∅ input stimuli on the Positive-Negative Output Task with global neuromodulation factor 0.5. Shading represents standard deviation. (*D*) Individual neuromodulators elicit unpredictable transforms of intracellular voltage traces in crustacean stomatogastric ganglion (STG) neurons LP and PD. Blue shaded traces indicate presence of neuromodulator serotonin. Reprinted and adapted from Neuron 76, Marder, Neuromodulation of Neuronal Circuits: Back to the Future, 1-11, 2012, with permission from Elsevier. (*E*) Five example neurons’ activity patterns from neuromodulated model RNN show complex nonlinear transformations analogous to crustacean STG activity changes under neuromodulation. Shading represents standard deviation. For (*A*), (*C*), and (*E*) arrowheads with dashed lines indicate timing of onset and offset of stimulus pulse (for + stimulus).

RNNs could successfully separate behaviors when recurrent weights only or recurrent, input, and output weights were all modulated at the same time, but not when only input or only output weights were modulated (Extended Data Fig. 4d–f). For the analyses presented below we focus on modulation of only the recurrent weights.

Application of this form of neuromodulation led to unpredictable transformations of individual neuron activity patterns and global network activity (Fig. 1e, Extended Data Fig. 2a,b). This is reminiscent of the observed neuromodulator effects on individual neuron activity patterns in the stomatogastric ganglion of crabs (Fig. 1d) and other organisms [15, 17, 18, 47], where neural activities shifts have been shown to be unpredictable and display non-linear transforms.

### Targeted neuromodulation can toggle behaviors across multiple global network states

In brains, neuromodulators are released in specific regions — some tightly localized, others broadcast widely — to influence local and global neural output. We found that RNNs with neuromodulated subpopulations of sizes across a broad range (100%–10% of the whole population) consistently supported the opposing behaviors of the task (Fig. 2a–c). Just as some neuromodulators affect neurons in a cell-type specific manner, for example selectively influencing activity of excitatory or inhibitory neurons with corresponding receptors [48], we found targeting of neuromodulator in this manner also was able to generate multiple behaviors (Fig. 2d,e).

**Figure 2.**
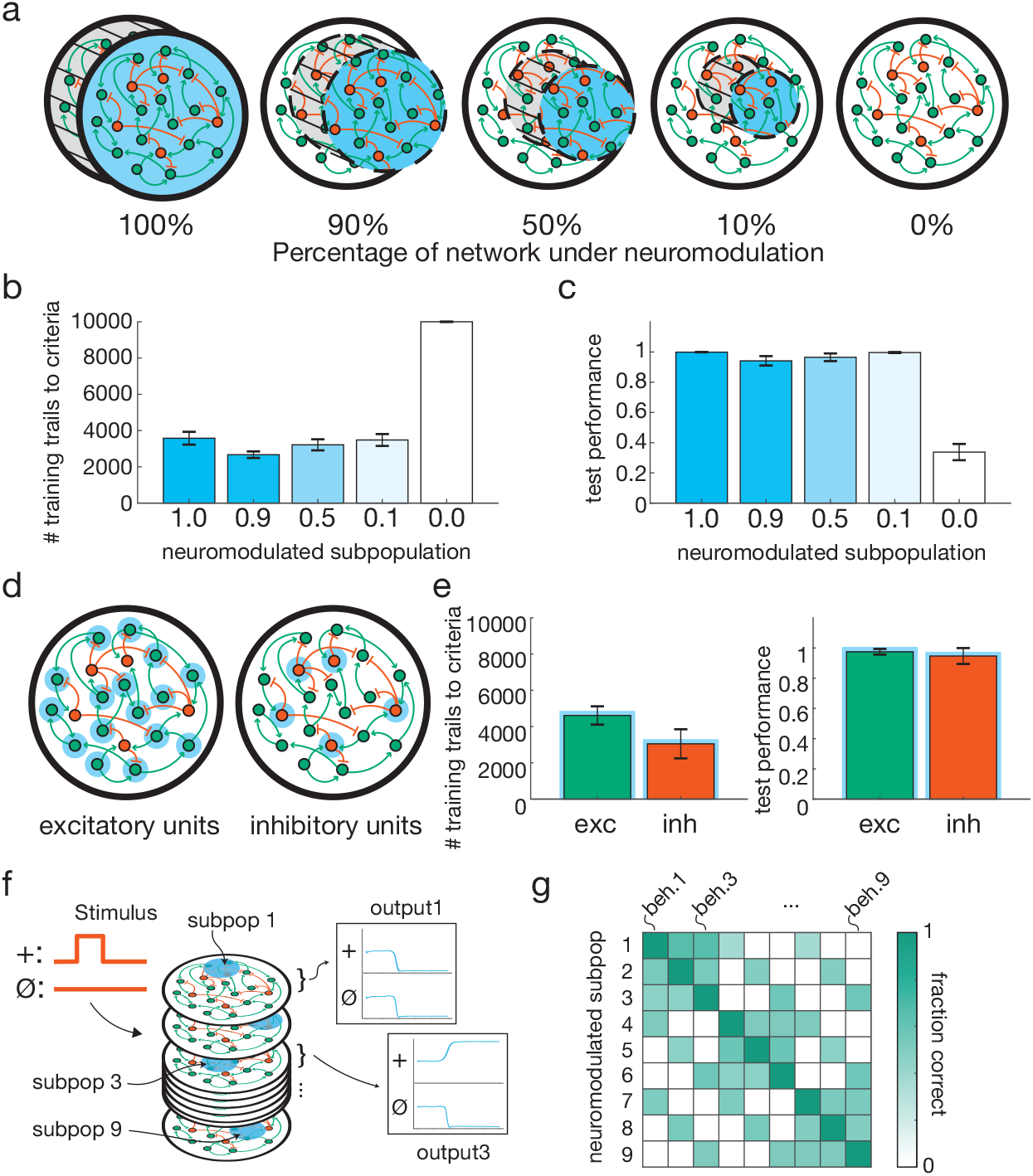
Targeted neuromodulation flexibly supports multiple behaviors. (*A*) A range of different sized neural subpopulations (0%-100% neural units) embedded within a RNN were neuromodulated (factor (*f*_*nm*_) = 0.5). 100% was positive control demonstrated in Fig. 1; 0% was negative control. (*B-C*) RNNs with embedded neuromodulated subpopulations across the size spectrum could support the behaviors of the Positive-Negative Output Task. (*B*) Number of training trials required to reach stop criteria (see Methods). (*C*) Test performance (1 on y-axis is 100% correct; see Methods). (*D*) Neuromodulation of exclusively excitatory or inhibitory neurons (blue annuli). (*E*) Excitatory or inhibitory neuromodulation supported learning of the Positive-Negative Output Task. (*F*) 9-behavior extended Positive-Negative Output Task with unique neuromodulated subpopulations (subpops) and example corresponding outputs. (*G*) RNNs successfully learned the task from (*F*) with 9 targeted subpops (each 10% of the RNN, non-overlapping; *f*_*nm*_=2.5). Application of neuromodulator to any subpop unlocked a specific behavior set (beh.) from the 9-behavior repertoire (fraction of trials correct is ≈1 on diagonal; see Methods). Off-diagonal fraction correct due to partial output overlap between behavioral sets (e.g. ∅ output as shown for output behaviors 1 and 3 shown in (*F*) is the same); for perfect performance, off-diagonals should be 0.5 or 0 depending on whether there is output overlap or not.

We next tested whether several unique behaviors could be learned and unlocked from a single network through targeted neuromodulation. To assess this, we created an extended version of the Positive-Negative Output Task in which the same two input stimuli (+ and ∅) could elicit 9 unique combinations of corresponding output behaviors from an RNN (see Methods for details). We found that neuromodulation of distinct subpopulations or with distinct neuromodulation levels could support the multiple output behaviors. In a single 200 unit neural network, we were able to generate 9 distinct behaviors by varying the location of where the neuromodulator effect was broadcast (Fig. 2f,g and Extended Data Figs. 5, 6).

### Distinct global network activity hyperchannels emerge from neuromodulation and exhibit non-linear transition dynamics

To understand how this form of neuromodulation leads to network behavior shifts, we analyzed the coordinated activity of all neurons in the RNN in the absence and presence of neuromodulator. At the individual neuron level, neuromodulation shifted the net difference of excitatory and inhibitory inputs, which in turn altered the recurrent propagation of activity over time and resultant internal network dynamics (Extended Data Figs. 7–9). At the whole population level, neural activity trajectories for the same stimulus with and without neuromodulator followed non-overlapping, stereotyped paths, or hyperchannels [49–51], in activity space (Fig. 3a). Activity hyperchannels refer to stereotyped trajectories in activity space (activity patterns) that define the constrained subspace of all possible network activities that is occupied when the network is in a given state and presented with a given stimulus [49]. Through amplification of synaptic weights, neuromodulation effectively resets all relative synaptic weights, altering activity flow patterns through the RNN (Extended Data Fig. 1f and 10a).

**Figure 3.**
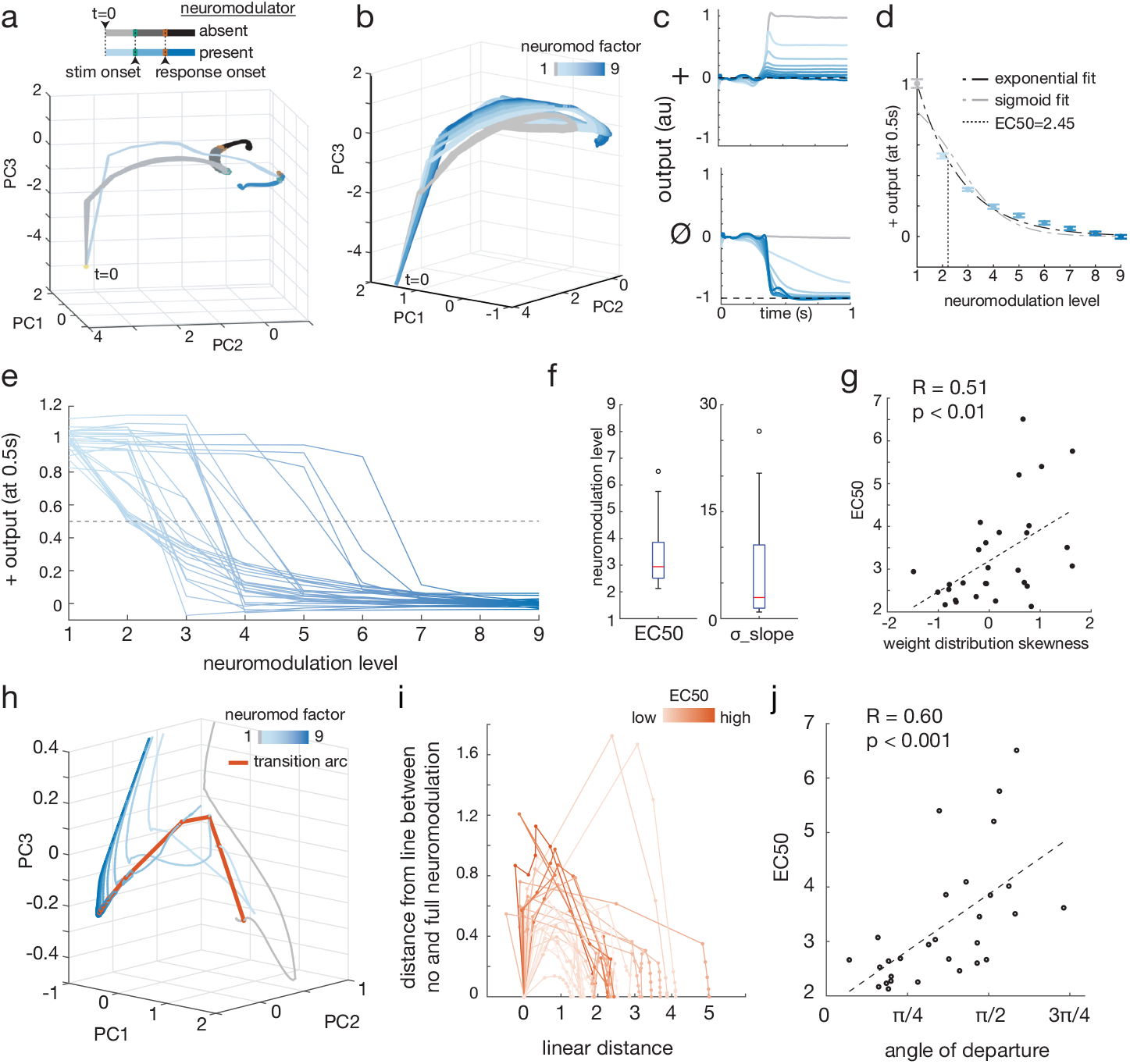
Neuromodulation separates activity hyperchannels with idiosyncratic nonlinear transition dynamics. (*A*) Global network activity dynamics after PCA-based dimensionality reduction for positive stimulus. Activity trajectories follow stereotyped, non-overlapping paths, or hyperchannels in activity space. (*B*) For a RNN trained with neuromodulatory factor of 9, intermediate levels of neuromodulation lead to partial transitions (traces in shades of blue) toward full neuromodulation activity hyperchannel (traces in darkest blue). (*C*) Partial neuromodulation maps to intermediate output behaviors. (*D*) Transition from none to full neuromodulation behavior is non-linear (best-fit exponential & sigmoid shown) and defines network sensitivity, measured as EC50. (*E*) Output transition for 29 networks independently trained on the Positive-Negative Output Task and tested across intermediate levels of neuromodulation. (*F*) 29 independently trained RNNs exhibit large variability in transition dynamics, with EC50 ranging from 2.1 to 6.5 and sigmoid slope (*σ* slope) from 0.9 to 26.3. (*G*) Network EC50 is positively and significantly correlated with skewness of global weight distribution. (*H*) Zoomed image demarcating path at a given timepoint over a network’s neuromodulation transition manifold (orange; transition arc) which connects hyperchannels corresponding to different neuromodulation levels in activity space (grey–blue trajectories). Blue shading same as in (*B*). (*I*) At a given timepoint (here t=100), intermediate levels of neuromodulation (represented by individual points) trace an arc between no neuromodulation (leftmost point on each curve) and full neuromodulation (rightmost point on each curve) states in phase space. Purely linear interpolation would lay along the x-axis at “distance from line (y-axis)” = 0. Each curve represents an individual network’s transition arc across neuromodulation levels. Arcs are shaded by relative EC50. Arcs represent cross-section of full transition manifold connecting hyperchannels across all timepoints at intermediate neuromodulation; each point on the arc is average of a cross section of a single hyperchannel. (*J*) Angle of departure (angle formed between direct path and first neuromodulation level hyperchannel) exhibits a strong positive correlation with network EC50. Transition in direction more orthogonal or away from full neuromodulation state results in lower sensitivity, i.e. higher EC50. For all PCA plots, top 3 PCs accounted for 80–92% of activity variance.

The distinct hyperchannels in activity space derive from a common underlying neural network. This suggests that there must be a transition between the hyperchannels that is accessible through intermediate amounts of neuromodulation. To characterize these transitional states, we applied intermediate levels of neuromodulation to the RNN after training, which mapped a smooth transition from trajectories of the non-neuromodulated hyperchannel to those of the fully neuromodulated hyperchannel (Fig. 3b). We found that intermediate neuromodulation generated intermediate outputs from the network (Fig. 3c). In this way, neuromodulated neural trajectories provide a means to dynamically respond to intermediate, even unexperienced neural states (e.g. hunger levels), just as in other work RNN neural trajectories have been shown to naturally address temporally varying sensory-motor patterns [49].

We then characterized how the RNNs’ output behaviors transitioned over the spectrum of applied intermediate neuromodulation levels. To compare different RNNs output behaviors, we took the output of each RNN at the midpoint of each trial for each level of neuromodulation, which captured each RNN’s output behavior response after any given stimulus presentation. We found that increasing neuromodulator levels led to non-linear progression from non-neuromodulated behavior to fully neuromodulated behavior (Fig. 3d; non-neuromodulated behavior in this simulation was output of +1; neuromodulated behavior was output of 0; behavior progression was best fit by exponential or sigmoid function).

We next assessed the degree of variability of this neuromodulator-dependent non-linear behavioral progression. It could be that similar neuromodulation of different networks results in similar transitions in behavior, representing a conserved property of neuromodulation transitions. Alternatively, individual differences in network connectivity could cause high variability in behavioral transition dynamics under neuromodulation. To evaluate this, we independently trained several RNNs generated with identical parameter constraints (n=29; see Methods for parameter constraints used). All RNNs exhibited non-linear transition dynamics best fit by an exponential or sigmoid function (Fig. 3e and Extended Data Fig. 10b), but networks’ sensitivities to neuromodulator and rates of transition varied drastically. To quantify this variability, we defined a “half maximal effective concentration” (EC50) as the amount of neuromodulator required to generate a half maximal output, analogous to the metric used to assess drug efficacy in pharmacology (see Methods). While all networks could produce intermediate outputs, the EC50 of individual networks trained with a full neuromodulator factor of 9 ranged from 2.1 to 6.5 (3.1x range; Fig. 3f, left) and rate of transitions (steepness of the transition dynamics sigmoid) varied widely as well from 0.9 to 26.3 (Fig. 3f, right). This result reveals a new, previously uncharacterized form of “circuit-based sensitivity” [52–55], that occurs even when all parameters of the circuit are selected in an identical manner and networks exhibit identical end behaviors. This variability, which is derived purely from the stochastic nature of weight initialization (the random starting configuration) and specific training experience (the specific sequence of instances experienced) may contribute to the wide individual variability of neuropsychiatric drug sensitivities observed clinically [20, 25, 56].

We next sought to characterize what properties of the networks contribute to the variability in sensitivity to neuromodulator. Since the neuromodulation effect we simulated acted on synaptic weights, we first analyzed individual network weight distributions. We found that the skewness of networks’ weight distributions exhibited a positive correlation with EC50s (R=0.51, p<0.01; Fig. 3g), suggesting networks with more positively skewed weights (longer tail of strong excitatory weights) were less sensitive to neuromodulator.

To further understand the source of network sensitivity variability, we characterized the shape of the networks’ activity transition curves across neuromodulation levels (Fig. 3h). At a given trial timepoint, purely linear interpolation yielded linear sensitivity relationships with invariant EC50 (Extended Data Fig. 11). In contrast, progressive neuromodulation defined an arc (Fig. 3h,i). Taken across all timepoints, this arc extended to form a curved transition manifold connecting each activity hyperchannel defined by a specific intermediate neuromodulation level. The geometry of this transition arc (measured as the angle of departure; see Methods) was strongly correlated to network sensitivity (Fig. 3j). This suggests that while individual networks achieve identical performance on the trained task (at no and full neuromodulation), the geometry of their population activities at intermediate neuromodulation levels is unique, leading to emergent sensitivity profiles. In particular, characterization of networks’ transition arcs revealed 1) transition dynamics are nonlinear due to curvature of the activity space transition, and 2) the specific shape of a network’s transition arc is tightly linked to its sensitivity to neuromodulator.

Excessive neuromodulation can also occur either pathologically or pharmacologically. To model this, we applied neuromodulation at levels higher than those used during training and found neural dynamics could sometimes (but not always) diverge from trained activity hyperchannels into an adjacent region of activity space, translating into inappropriate output behavior (Extended Data Fig. 10c–e).

### Neuromodulated RNN replicates dopamine-mediated starvation-dependent sugar sensitivity in *Drosophila*

Given our finding that neuromodulation provides a natural means of handling intermediate, unexperienced neural states, we next sought to evaluate our model’s ability to recapitulate the behavioral effects of neuromodulation observed *in vivo*. The neuromodulator dopamine controls the sugar sensitivity behavior of *Drosophila* [1], as measured by proboscis extension reflex (PER) probability, which increased with both duration of starvation (fed, 1 day, 2 day starved) and concentration of L-dopa administered in their diet (0, 3, 5 mg/ml) (Fig. 4a). PER is a behavior of *Drosophila* in which they extend their proboscis in response to a desirable food stimulus.

**Figure 4.**
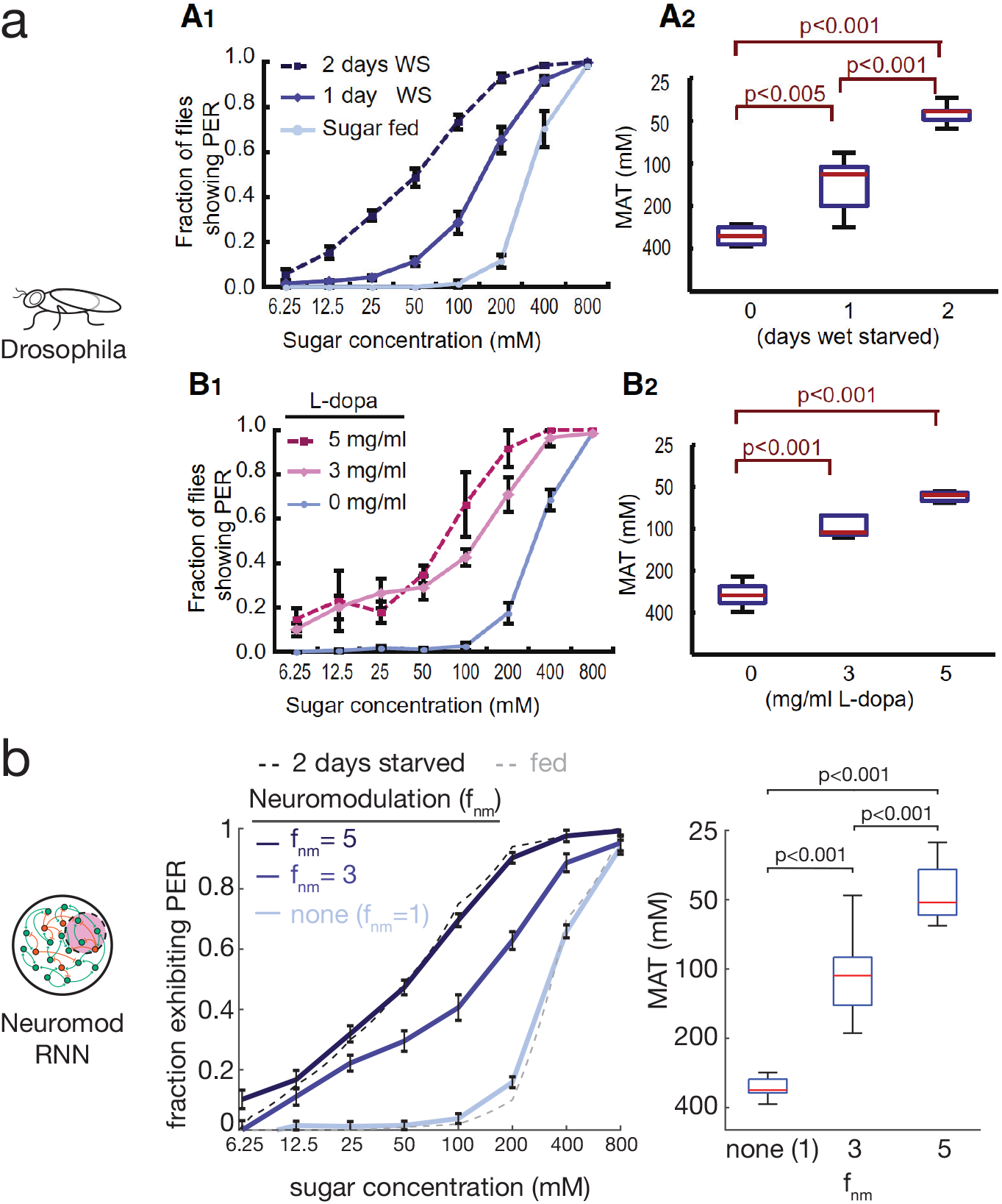
Neuromodulated RNNs reproduce *Drosophila* sugar sensitivity behaviors. (*A*) *Drosophila* sugar sensitivity behaviors from Inagaki et al. 2012 measured as proboscis extension reflex (PER) behavior vs sugar concentration and mean acceptance threshold (MAT; the sugar concentration that elicits 0.5 PER fraction). Reprinted from Cell 148, Inagaki et al., Visualizing Neuromodulation In Vivo: TANGO-Mapping of Dopamine Signaling Reveals Appetite Control of Sugar Sensing, 583-595, 2012, with permission from Elsevier. (*B*) Neuromodulated RNNs trained on extremes of *Drosophila* sugar sensitivity (no neuromodulation factor (*f*_*nm*_=1) for fed and *f*_*nm*_=5 for 2 days starved) exhibit similar intermediate (*f*_*nm*_=3; untrained) and extreme (no neuromodulation (*f*_*nm*_=1) and *f*_*nm*_=5; trained) behaviors (n=10; error bars are SEM; same statistical test as in Inagaki et al. 2012 for boxplots, see Methods).

To assess if our neuromodulation model could reproduce these results, we trained RNNs with neuromodulated subpopulations (20% subpopulation) to reproduce the fed and 2 day starved sugar sensitivity curves of flies (no neuromodulation (*f*_*nm*_=1) for fed; *f*_*nm*_=5 for 2 day starved). We then tested the RNNs’ behaviors at an intermediate, never-before experienced neuromodulator level (*f*_*nm*_=3). The RNNs produced a shifted sensitivity curve very similar to that exhibited by 1 day starved flies and the flies fed an intermediate L-dopa concentration of 3 mg/ml (Fig. 4b), as measured by sugar sensitivity curve shift and mean acceptance threshold (MAT), which is the sugar concentration at which 50% of flies or networks exhibit the PER (see Methods).

The behavior of the RNNs reliably mimicked the intermediate behaviors of flies *in vivo* because intermediate neuromodulation caused a continuous shift in the RNN’s activity hyperchannel between “fed” and “2 day starved/5 mg/ml L-dopa” hyperchannels (Extended Data Fig. 12a–c). Furthermore, our model reveals that transition manifolds (defined by the hyperchannels connecting intermediate neuromodulation levels) are unique to networks, predicting wide-ranging natural variability in fly starvation-based sugar sensitivity profiles (Extended Data Fig. 12d). Neuromodulation in our model leads to natural handling of never-before experienced neural states by creating a network configuration such that the neuromodulatory transition manifold has a geometry that leads to intermediate outputs. Although the dopamine-dependent control of sugar sensitivity in *Drosophila* is likely regulated by other effects as well, our model shows that single scalar synaptic scaling is already sufficient to fully account for the starvation-dependent sugar sensitivity shifts observed.

### Electrical modulation shifts neural dynamics through an independent circuit effect

Other endogenous and exogenous influences can alter neural circuit dynamics through mechanisms that may be shared or independent, and understanding relationships between such interventions is vital for safe and effective treatment. We used our model to compare whether exogenously delivered electrical modulation of a neuromodulated circuit — analogous to use of optogenetics experimentally [57] and deep-brain stimulation (DBS), transcranial magnetic stimulation (TMS), or transcranial direct current stimulation (tDCS) clinically [58–61] — alters network activity in an analogous manner to chemical neuromodulation or operates through an independent effect (Fig. 5a).

**Figure 5.**
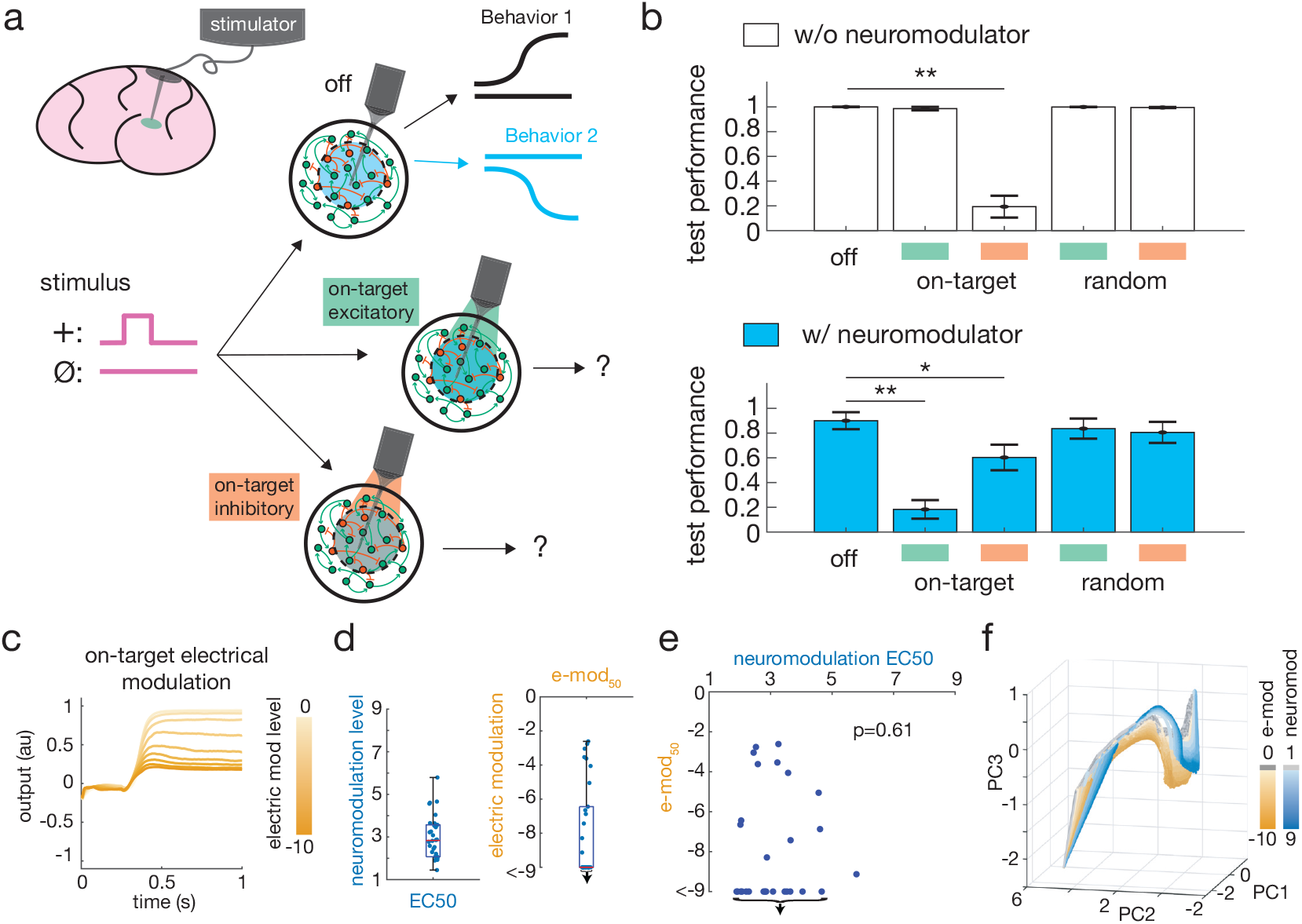
Targeted electrical modulation shifts network dynamics through independent circuit effect. (*A*) Schematic of DBS and analogous electrical modulation (e-mod) of a neuromodulated RNN on the Positive-Negative Output Task. (*B*) Test performance was significantly impaired in absence of neuromodulation (white bars) when inhibitory on-target electrical modulation (e-mod) was given (p=3.91e-11) and with neuromodulation (blue bars) when excitatory and inhibitory on-target e-mod was given (p=2.20e-08 and p=2.18e-02, respectively). *:p<0.05, **:p<0.01, Student’s *t* -test. Green indicates applied excitatory e-mod; orange indicates inhibitory e-mod. Performance decreased due to behavioral shifting directionally toward opposing neuromodulation condition (see Extended Data Fig. 13a,b). Error bars are SEM over 10 independent networks (see Methods). (*C*) Example RNN output in absence of neuromodulation to + stimulus with increasing inhibitory e-mod. (*D*) For 30 RNNs: Left: neuromodulation EC50. Right: electrical e-mod_50_. Down arrows in (*D-E*) indicate RNNs that did not achieve e-mod_50_ at maximum stimulation. (*E*) No significant correlation between networks’ EC50 and e-mod_50_. (*F*) Neuromodulation and electrical modulation push activity trajectories along different transition manifolds enabling independent transition dynamics.

We simulated electrical modulation as added excitatory or inhibitory current (see Methods) to RNNs that had been trained with neuromodulated subpopulation (50% of neurons) on the Positive-Negative Output Task. We found that while electrical current given to random subpopulations did not affect the RNNs’ performances, current delivered to the neuromodulated subpopulation could significantly affect network output (Fig. 5b), shifting behaviors directionally toward the opposing neuromodulation condition behavioral set (Extended Data Fig. 13a,b).

In each RNN, increasing the level of targeted electrical input — for example inhibitory modulation in the absence of neuromodulator — led to a graded transition in behavior similar to the transition observed with graded administration of neuromodulator (Fig. 5c ; see Methods). To compare these conditions, we measured the amount of electrical input that led to output of 50% of the fully-neuromodulated condition (e-mod_50_) analogous to EC50 for neuromodulator levels. Interestingly, RNNs also exhibited idiosyncratic circuit-based sensitivity to electrical modulation, with e-mod_50_ under inhibitory modulation ranging from -2.6 input current to not achieving e-mod_50_ by the maximum modulation we administered (−9 input current; >3.5x range) (Fig. 5d, right).

We then assessed whether networks’ electrical and chemical sensitivities were related, and found no significant correlation (Fig. 5e). Since some RNNs’ outputs did not reach e-mod_50_, saturating before the maximum electrical input given, — further evidence of a different mechanism at play — we also measured each RNN’s output at maximum electrical input (input=-9) and similarly found no significant correlation to EC50 (Extended Data Fig. 13c). The lack of correlation suggests that networks insensitive to chemical modulation may still be highly sensitive to electrical modulation and vice versa. Consistent with this, we found that electrical modulation progressively shifted population dynamics along a manifold transition distinct to neuromodulation, enabling different rates of transition (Fig. 5f). This may help explain why some patients who fail pharmacologic treatments sometimes respond dramatically to electrical neuromodulation interventions like DBS or TMS. In addition to their localized effects to specific brain regions, DBS and TMS utilize an different mechanism to chemical modulation, leveraging an independent circuit-sensitivity to achieve therapeutic efficacy.

## Discussion

Neuromodulation in brains drives unique neural function in health and disease. Using an RNN model, we showed how neuromodulators can act through simple scaling of synaptic weights to generate unique behavioral modes from a single RNN. Our model sheds light on how neuromodulation works biologically, specifically neuromodulator effects on synaptic weights [15, 17, 18, 27, 28, 38]. Using a series of tractable tasks, we showed how this unique and previously uncharacterized control mechanism in RNNs is able to rapidly reconfigure a network with only a single scalar input.

We make two specific predictions. First, we posit that neural networks exhibit idiosyncratic transition profiles resulting in varying sensitivities to neuromodulation. Networks’ sensitivities depend on the geometry of their transition profiles and weight distributions, specifically the shape of the transition manifold defined in activity space. Second we predict that chemical neuromodulation acts on networks via a separate transition manifold to that of electrical stimulation, resulting in uncorrelated sensitivity profiles between these different modes of neuromodulation.

These predictions can be directly tested. Although there is already indirect evidence that neural networks in the brain have idiosyncratic sensitivities to neuromodulation via variable response to treatments like SSRIs, direct evidence could be gained via experiments applying varying amounts of neuromodulator (e.g. concentrations of serotonin) to a specific circuit, measuring network output, and repeating this over several circuits (*in vitro*) or organisms (*in vivo*) to compare their response profiles. Neural recordings of the circuits during these experiments would enable assessment of transition profiles in neural activity space to test our hypothesis about transition manifold geometries.

At a behavioral level, our simulations studying *Drosophila* behavior could be tested experimentally by measuring the behavioral transitions of individual flies and measuring the variability compared to our model’s prediction. Our second prediction could be tested with experiments applying both chemical neuromodulation (e.g. application of serotonin to a circuit) as well as direct electrical stimulation (via electrode or optogenetics). By varying the degree of each mode of neuromodulation, output response profiles can be measured and compared to characterize if each mode of neuromodulation operates along an independent, uncorrelated transition profile as predicted by our simulations. Here again, simultaneous neural recording would enable activity space characterization and comparison to our model’s transition manifold predictions.

Our model provides insights into neuromodulation that are related to other recently elucidated principles of neural computation. We showed that neuromodulation leads to separation of distinct activity hyperchannels, similar to those observed by Goudar and Buonomano [49] and Nieh et al. [51], with neuromodulation effectively disentangling neural trajectories by separating them in phase space analogous to the work of Russo et al. [62] in motor cortex. Just as neural trajectories provide transformations in phase space that naturally handle temporal variation of sensory-motor patterns [49], neuromodulation leads to transformations in phase space that elucidate a biological mechanism for handling intermediate and continuously transitioning neural states, even if never experienced before. We demonstrated the biological use of this property through replication of Inagaki et al.’s findings in *Drosophila* [1]. In this way, the level of neuromodulation acted as a controller on the amount of disentangling of neural trajectories, using internal neural state (amount of weight modulation) to control output behavior. Such a system is robust to external noise (Extended Data Fig. 1), since far apart neural states generate trajectories that are widely separated in phase space.

Through this analysis we discovered a type of “circuit-based sensitivity” previously unreported to our knowledge. Previous studies have characterized network parameters (e.g. neuron types [54], maximal gain conductances [52, 55, 63], specific rectifying currents [53]) that can alter sensitivity of a network when varied and demonstrate that individual networks can behave differently. Here, we identify a form a “circuit-based sensitivity” that arises when networks are created under exactly the same conditions (no network parameter is systematically varied between them) and generate exactly the same end behaviors. The variability in sensitivity emerges only in their transition dynamics and results purely from the stochastic nature of network weight initialization and unique individual experiences (random sequences of the exact same training data). This type of “circuit-based sensitivity” may contribute (along with other forms) to the high variability in drug and other therapeutic response observed clinically [20, 25, 56], alongside more standard explanations like enzyme variant-dependent drug metabolism and clearance rates [24, 26, 64, 65].

This emergent sensitivity property of neuromodulated networks is related to the well-known “many solutions” phenomenon where different weight configurations can produce identical output [66–68].

Our model demonstrates a new facet of circuit-variability derived from the transition dynamics in RNNs under varying neuromodulation. We reveal that unique geometric configurations of phase space underlie emergent network sensitivity profiles. An interesting future research question is whether the sensitivity described in this study is related to other forms of circuit sensitivity previously described, for example, the variability observed in 2-16 neuron models comprised of Hodgkin-Huxley-type neurons initialized with different maximal gain conductances. Are particular weight profiles in the larger networks analogous to a range of maximal conductance parameters as was investigated in these smaller network models [55], perhaps also relating to “sensitive” and “insensitive” directions both in parameter space [63] and activity space?

Future studies aimed at identifying circuit parameters that control neural activity transition dynamic geometry will be critical for understanding and use in therapeutic optimization. New insights into network sensitivity will be gained by developing useful metrics to characterize how a given network’s activity space is arranged (the shape of the transition surface) and understanding the relationship between networks’ synaptic weight profiles and their activity space geometries. Our study begins to contribute to this effort by identifying the activity transition surface geometry as a critical factor in predicting network sensitivity. The idiosyncratic nature of circuit-based sensitivity aligns with current efforts in precision medicine calling for the need to consider each patient as an idiosyncratic individual — here we provide computational evidence for this claim and its particular importance in neuropsychiatric treatment [69]. Fully understanding the relationship between chemical and electrical modulation and sensitivity is also crucial. Although our simplified model suggests how the modes of modulation influence dynamics (see Extended Data Appendix B), further analytical and experimental investigation into their relationship as network dynamics evolve over time could provide deeper insights.

Our formulation of the neuromodulatory effect on synaptic weights is a simplification of the true biological mechanism. Elaborating our model to support differential weight modulation (e.g. via multiple neuromodulators and neuromodulator receptor subtypes on specific cell-types) [4, 70, 71], neuromodulator multiplexing [44, 48], and metamodulation [6, 28, 72] will likely lead to even more sophisticated network behavior. Our model also uses arbitrary neuromodulation levels, whereas brains likely use specific levels for optimal functionality. Future investigation into which levels are optimal and methods of learning these will be important.

Additionally, the tasks we used to investigate how this aspect of neuromodulation works were relatively abstract and simple. This allowed us to tractably demonstrate and probe properties of neuromodulation and how they affect network computation and behavior. Future investigation into whether this simple mechanism alone is sufficient to support libraries of much more complex behaviors within a network, like those exhibited by humans in everyday real life and when more sophisticated versions like multiplexing and metamodulation are required is a fascinating direction of future research.

Our model of neuromodulation suggests several interesting connections to biological and clinical phenomena, but also has limitations. In addition to those noted above due to the simplified model we investigated, our study is rooted in artificial neural network (ANN) models. This approach has major advantages in studying fundamental principles of how neuromodulation of synaptic weights can modify parallel, distributed circuit computations, but also is strongly limited in its explanatory power of the full biological phenomena, as in psychiatric diseases, which have many more relevant parameters (neuron subtypes, circuit architecture, glia, environmental and historical factors, vasculature, etc.) not accounted for in our ANN models. Given these limitations of our model, our findings require experimental validation.

Finally, neuromodulation provides interesting directions for machine learning (ML). By separating synaptic memory regimes in a single network, we demonstrate how a network can have much greater flexibility and increased capacity, supporting a library of unique behaviors for overlapping external contingencies. Furthermore, each behavior can be rapidly accessed through targeted application of the relevant neuromodulatory factor. High capacity, compact networks with high-speed access to different output modes presents a promising component for ML development and storage-limited applications like edge computing. Additionally, through the separation of memory regimes that effectively splits a single RNN into multiple processors, this mechanism may provide a means of realizing the super-Turing capability of specific RNN configurations as defined by Cabessa and Siegelmann [73]. Future theoretical assessment of neuromodulated RNNs’ capacity will establish if this simple mechanism is sufficient to exceed the Turing limit.

## Methods

### Positive-Negative Output Task

In the Positive-Negative Output Task two possible stimuli could be given to the agent (RNN in our simulations): either a positive pulse referred to as positive stimulus or +; or no pulse, also referred to as null stimulus or ∅ (See *Supplementary Information* for details of input signals). The agent is trained to give a positive output (+1) for the positive stimulus and zero output (0) for the null stimulus. When a neuromodulation effect is applied (e.g. broadcast synaptic scaling by factor of 0.5), the agent is trained to output a different set of behaviors to the same stimuli: zero output for the positive stimulus and negative output for the null stimulus.

The Positive-Negative Output Task has many similarities to the well-known Go-NoGo task which is a widely used experimental paradigm. The Positive-Negative Output Task differs in that there are two behavioral repertoires which can be elicited depending on the internal state of the agent which corresponds to the presence or absence of the neuromodulatory effect in our RNN models. Internal state typically transitions over longer timescales than stimulus-driven shifts in behavior; our task was designed to depict these longer timescale shifts in internal state that drive diverse behaviors.

The 3 behavior and 9 behavior variants of the Positive-Negative Output Task followed a similar paradigm with additional added behaviors. In the 3 behavior version, a third behavior of positive stimulus → negative output, null stimulus → positive output was added. The 9 behavior version was similarly elaborated with 9 possible behaviors. See *Supplementary Information* for details. In our simulations each trial lasted 200 timesteps of 5ms each for a total represented duration of 1 second.

### Neuromodulatory neural network model

For our simulations we used a continuous rate recurrent neural network (RNN) model with biologically plausible parameters similar to RNNs in prior works [74, 75]. Consistent with biological neural networks, we implemented Dale’s Law using the method in Song et al. [76] such that each neuron was either excitatory or inhibitory. For all our simulations we used a RNN with *N* = 200 neuron units, 80% excitatory and 20% inhibitory. In the RNN each neuron can be connected to any other neuron with probability *p*_*con*_ (*p*_*con*_ was initialized at 0.8 in our simulations), and each neuron receives weighted input from connected neurons to produce a firing rate governed by the neural dynamical equation

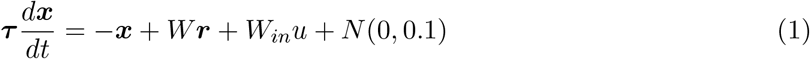

where ***τ*** ∈ *R*^1×*N*^ is the synaptic decay time constant for the *N* neurons in the network, ***x*** ∈ *R*^1×*N*^ is the synaptic current variable for the *N* neurons, *W* ∈ *R*^*N*×*N*^ is the matrix of synaptic weights between all *N* neurons, ***r*** ∈ *R*^1×*N*^ is the output firing rates of the N neurons in the network, *W*_*in*_ ∈ *R*^1×*N*^ are weights associated with external input *u*, and *N* (0, 0.1) is added noise drawn from a normal distribution with mean 0 and variance 0.1. The output firing rate for the neurons is given by an elementwise nonlinear transfer function transformation of the synaptic current variable. In our network we used the standard logistic sigmoid function as implemented by prior models [74, 77]:

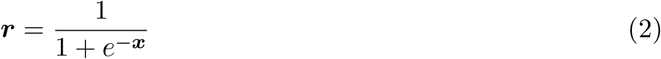

The synaptic connectivity matrix *W* was randomly initialized from a normal distribution with zero mean and standard deviation 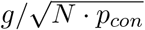, where *g* is the gain. We set *g* = 1.5 as previous studies have shown that networks operating in a high gain regime (*g ≥* 1.5) support rich dynamics analogous to those of biological networks [74, 77, 78]. The synaptic decay time constants were randomly initialized to a value in the biologically plausible range of 20–100 ms. As in Kim et al. 2019, we used the first-order Euler approximation method to discretize equation (1) for the simulations; for neuron *i*:

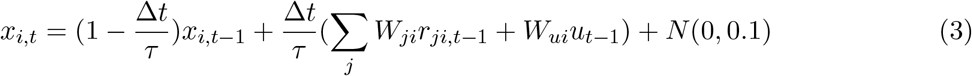

Output was generated by taking all recurrent network neurons’ activities and passing them through a weighted output unit

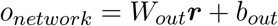

where *W*_*out*_ ∈ *R*^*N*×1^ are the neural output weights and *b*_*out*_ is the output unit’s bias term. RNNs were trained by backpropagation through time using AdamOptimizer with a least root squared error objective function:

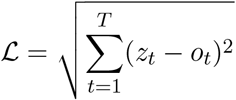

where *z*_*t*_ is the desired output and *o*_*t*_ is the network output at time *t* in a trial of total length *T* (*T* = 200 timesteps where each timestep represents 5 ms, for a total trial length of 1 second in our simulations). For the Positive-Negative Output Task, desired output was a 200 timestep output trace as described above under “Positive-Negative Output Task.” For the Drosophila task, the desired output was a single target value as described below under “*Drosophila* sugar sensitivity task;” in this case only one timestep of output (the final network output at t=100) contributed to the loss.

To apply an amplifying or dampening neuromodulatory effect, target neurons’ weights were scaled by the neuromodulatory factor. For whole network neuromodulation this effect was applied to all neurons in the RNN; for subpopulation neuromodulation the effect was applied only to the selected subpopulation of neurons.

For the Positive-Negative Output Task, RNNs were trained until one of two possible stop criteria was met: 1) average trial root square error over the last 50 trials was under a threshold of 1, or 2) 10,000 training trials was reached. Performance on the task was then assessed by evaluating the percentage of test trials that matched the following performance criteria: for positive output trials, output was required to reach 1.0 ± 0.2 by timestep 120 (full trial was 200 timesteps); for zero output trails, output was required to be 0.0 ± 0.2 and for negative output trails −1.0 ± 0.2 at timestep 120.

For the Drosophila task, RNNs were trained until one of two possible stop criteria was met: 1) average trial square error over the last 100 trials was under a threshold of 0.005, or 2) 10,000 training trials was reached. Output of RNNs was then assessed to determine network MAT (see Mean acceptance threshold (MAT) description below) and behavior at inputs corresponding to standard sugar concentrations taken from Inagaki et al. 2012 [1].

### Comparison to context-dependent cued model

For comparison, we created a cue-driven RNN model and trained it on the Positive-Negative Output Task. The cue, which we refer to as the “context cue”, was delivered through an additional input hyperchannel and signaled which output behavior was desired. We created models with two types of cues: transient cues were constant value inputs present only during the stimulus input period (t=1 to t=75); persistent cues were constant value inputs across the whole trial. For comparisons between models, we ran models with cue pairs of +1.0/-1.0 (2.0 sep in Extended Data Fig. 1d,e), +0.5/-0.5 (1.0 sep), and +0.2/-0.2 (0.4 sep).

We compared neuromodulatory and context-cued RNNs tolerance to noise in input signals and noise in neuromodulatory signal. For each noise condition we tested, we ran 100 simulations in 4 replicate RNNs and measured consistency of final network states using a measure of mean Euclidean distance. For details of noise simulations comparison to context-dependent cued model see *Supplementary Information*.

### Neuromodulation of multiple subpopulations and multiple levels

For neuromodulation of non-overlapping subpopulations, same-sized groups of neurons were choosen randomly without any overlap and neuromodulator applied to each for a given behavior. For overlapping subpopulations, groups of neurons were chosen randomly allowing overlap (Extended Data Fig. 6).

Neuromodulation at different levels (“multi-factor networks”) was done by applying different neuromodulation factors (*f*_*nm*_). For the 9-behavior Positive-Negative Output Task this was done using factors ∈ [1 : 1 : 9], i.e., for Behavior 1 no factor was applied (*f*_*nm*_ = 1), for Behavior 2 *f*_*nm*_ = 2, for Behavior 3 *f*_*nm*_ = 3, et cetera.

To test networks across the range of subpopulation sizes with single or multiple neuromodulator factors on n-behavior Positive-Negative Output Tasks, stop criteria were adjusted to account for the increased behaviors: 1) average trial root square error over last n*25 trials was under threshold of 1, or 2) 15,000 training trials was reached. Performance on the tasks was assessed as before. For overlapping subpopulations, overlap was quantified in two ways. For each network, the number of neurons neuromodulated in 2 or more subpopulations was measured (Extended Data Fig. 6d). Overlap was also quantified by measuring the average number of neuromodulated subpopulations a neuron in the network was a member of (Extended Data Fig. 6e).

### Single neuron inputs, functional clustering, and selectivity index

Neural activity in a RNN is a complex function of all the neuron activities tracing all the way back in time. To understand how neuromodulation shifted synaptic inputs at the single neuron level, we considered the first time point in a trial. For any trial, at t=0 all activities are randomly initialized from a normal distribution. As a result, at t=1, a neuron reacts only to the weighted inputs of its incoming connections, uncontaminated by propagating recurrent activity dynamics from past timepoints. Analysis of neuron activity at this timepoint is shown in Extended Data Fig. 7a–c.

In order to examine whether trained models contained functionally specialized neurons, we grouped neuron activities by combination of subtask (which maps one-to-one with neuromodulatory state) and stimulus given (“stimulus-subtask combinations”). We averaged activity of each neuron over time and trials within each group. This resulted in a matrix of time-trial averaged neuron activities with a number of rows equal to the stimulus-subtask combinations, and number of columns equal to the number of neurons. Using k-means, we clustered neurons with similar activity levels across stimulus-subtask combinations. We computed a silhouette score to find the optimal number of clusters, which for the RNN in Extended Data Fig. 7, was 6. The silhouette score computed for 5 and 6 clusters differed only by 0.7% and the additional cluster was very small and similar to an existing cluster, so we conducted further analysis with 5 clusters for simplicity.

To measure the selectivity of individual neurons for particular stimulus-subtask combinations, we calculated a “selectivity index” (si) for each neuron j:

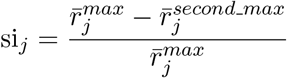

where 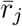 is the average firing rate of neuron j over the trial duration, 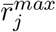 indicates the maximum 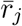 across all the stimulus-subtask combinations and 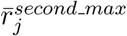 indicates the second highest 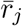 over all the stimulus-subtask combinations. The selectivity index thus captures a normalized approximation of how uniquely active a neuron was for a given stimulus-subtask combination.

### Network population dynamics

To represent whole network population activity dynamics we sought a low dimensional representation of whole population activity. We used principal component analysis (PCA) since the leading components capture the largest projections of activity variability, which we hypothesized would effectively separate our neuromodulatory conditions if large differences occurred [79]. We found this was the case. We found qualitatively similar results using multidimensional scaling which finds projections designed to best preserve distances in high-dimensional activity space. For our figures we display the first 3 PCs, as these captured a large amount of the activity variance (80–92% explained across the analyses) and effectively represent the activity dynamic differences in the analyses.

To map the neuromodulation-dependent activity subspace, we generated 100 independent stimuli series consisting of random numbers drawn from a uniform distribution between 0 and 1 at each time point (t=0 to t=200) and fed this into the RNN with and without neuromodulation (shotgun stimulus mapping from Extended Data Fig. 10a), which defined non-overlapping subspaces of activity space.

To analyze neuromodulation transition curves we compared activity under intermediate neuromodulation levels with linear interpolation. Linear interpolation was done by evenly dividing the distance between no and full neuromodulation activity states into 9 sections analogous to the 9 neuromodulation levels assessed. These 9 points in activity space were then used to generate output that is plotted in Extended Data Fig. 11. The geometry of the neuromodulation-based transition was assessed by calculating the Euclidean distance of intermediate neuromodulation level states at a given trial timepoint to the nearest point on the line connecting no and full neuromodulation states at that timepoint. These distances are plotted in Fig 3i. The angle of departure (AoD) was defined as the angle formed by the line between no and full neuromodulation states and the line between no neuromodulation and the first neuromodulation level states, which can be calculated as:

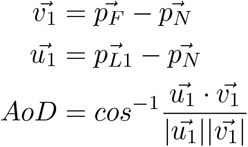

where 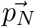 is the network state with no neuromodulation, 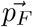 is the network state with full neuromodulation, 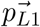 is the network state with the first level of neuromodulation.

### EC50

The EC50 of a network was defined as the level of neuromodulation that led to half the output of full neuromodulation, analogous to EC50 measured in dose-response curves for pharmaceuticals: the dose (“effective concentration”) that drives a half-maximal (50%) response. For the results reported, we used EC50 calculated for the positive stimulus. For this stimulus in the Positive-Negative Output Task, a non-neuromodulated network outputs +1 and a fully neuromodulated network outputs 0 (measurements for output level were taken at 0.5 s through the trial); the EC50 for the network in this case is the amount of neuromodulator required to output 0.5. The EC50 was calculated by fitting a sigmoid curve to the progression of output (from +1 to 0 in this case) with increasing neuromodulation level (Fig. 3d)

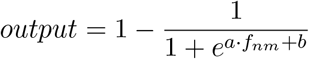

where *f*_*nm*_ is the neuromodulation level. EC50 neuromodulation level was calculated by finding the intersection of the sigmoid and the half-maximal output; for half-maximal output of 0.5, EC50 = −*b/a*. Sigmoid curves were fit using a least squares fit.

To assess the variability of individual network sensitivities to intermediate levels of neuromodulation, 29 RNNs were independently trained with a neuromodulation factor of 9. Since our focus in this analysis was on the variability of sensitivity to neuromodulator exhibited by networks, *∼*30 independent samples of RNNs was sufficient to show the high variability — a 3.1x spread, ranging from 2.1 to 6.5 with a full neuromodulation factor of 9. This demonstrated that occurrence of low and high sensitivity networks occurred with relatively high likelihood; roughly at least 1/30 or 3%.

### *Drosophila* sugar sensitivity task

We implemented a computational version of Inagaki et al. 2012 to train our network models. During training, models were presented with a constant sugar concentration (external input proportional to sugar concentration) for 100 timesteps (equivalent to 500 ms) and trained to output a probability of proboscis extension reflex (PER), a behavior exhibited by *Drosophila* where the proboscis is extended in response to a desirable food stimulus. This network output represents the upstream signal of executing the PER behavior; thus when a network output was 50%, 5 times out of 10 that network would do the PER. This directly would translate into the fraction of flies on average that would exhibit PER in that condition as measured in Inagaki et al. 2012. Curves in Fig. 4b show average behavior of multiple networks, where each network showed unique behavior transitions as reflected in Extended Data Fig. 12d. For fed and 2-day starved training we used a piece-wise linear approximation estimated from Inagaki et al. 2012 to determine desired output values. To compare boxplots of MAT, one-way ANOVA followed by *t* -test with Bonferroni correction was used as in Inagaki et al. 2012.

During training, networks received an input stimulus corresponding to a sugar concentration drawn from a continuous uniform distribution between a maximum and minimum sugar concentration (6.25mM and 800mM as in Inagaki et al. 2012 [1]). Networks were trained to output the sugar sensitivity curves corresponding to fed (no neuromodulation) and 2-day starved (full neuromodulation, factor 5). After training, networks were tested with no neuromodulation, full neuromodulation (factor 5), and an intermediate neuromodulation level not seen during training (factor 3).

### Mean acceptance threshold (MAT)

MAT is the sugar concentration at which *PER* = 0.5. Analogous to the analysis done for flies in Inagaki et al. 2012, for each RNN a sigmoid was fit

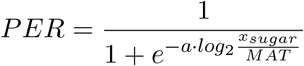

where *a* is the slope of the sigmoid. When *PER* = 0.5 then *x*_*sugar*_ = *MAT*. Sigmoid curves were fit using a least squares fit.

For intermediate neuromodulatory level (*f*_*nm*_=3) MAT variability analysis, a normalized change in MAT (%ΔMAT) was calculated:

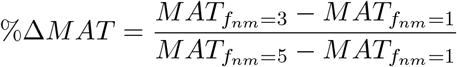

where 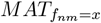 is the RNN’s MAT with neuromodulation factor *x*. %ΔMAT gives a network normalized metric for how much the intermediate neuromodulation (*f*_*nm*_=3) moved the fly from no neuromodulation (*f*_*nm*_=1) to full neuromodulation (*f*_*nm*_=5) sensitivity.

### Electrical modulation

We administered electrical modulation as an external current applied for the duration of the trial. “On-target” modulation was applied to the neuromodulated neuron population and “random” modulation was applied to a randomly selected group of neurons of equal size; these could include both neuromodulated or non-neuromodulated neurons. All neurons (both excitatory and inhibitory) within the selected subpopulation were given identical external current modulation. For fixed electrical modulation simulations (Fig. 5b and Extended Data Fig. 13a,b), a current of magnitude 1 was applied (+1 for excitatory modulation; -1 for inhibitory).

The initial experiment testing the e-mod effect (Fig. 5b and Extended Data Fig. 13a,b) was done in 10 independently trained RNNs with 50% subpopulation of neurons neuromodulated by a factor of 0.5. After identifying the effect of e-mod on RNNs, we sought to compare e-mod sensitivity to neuromodulation sensitivity. To do this, we trained 30 RNNs with 50% subpopulation neuromodulated and a factor of 9 (similar to the sensitivity analysis in Fig. 3). We then administered graded electrical modulation (e-mod ∈ −{0.0, 0.5, 1.0, 1.5, 2.0, 2.5, 3.0, 3.5, 4.0, 5.0, 6.0, 7.0, 8.0, 9.0}) and assessed network sensitivity (e-mod50) the same way as for neuromodulation (EC50). For these network we also measured neuromodulation sensitivity for comparison. These results are shown in Fig. 5c–f and Extended Data Fig. 13c. For details see *Supplementary Information*.

Test performance (Fig. 5b and Extended Data Fig. 13a,b) was measured in the same way as described previously for the Positive-Negative Output Task by assessing the percentage of test trials that matched the following performance criteria: for positive output trials, output was required to reach 1.0 ± 0.2 by timestep 120 (full trial was 200 timesteps); for zero output trials, output was required to be 0.0±0.2 and for negative output trails −1.0±0.2 at timestep 120. To assess decreases in test performance shown in Fig. 5b, further analysis was to done characterize the breakdown of specific behaviors as shown in Extended Data Fig. 13a,b.

Though similar to exogenous cues [9] in the mode of delivery (compare *W*_*ui*_*u* and *u*_*Estim*_ terms in last equation of Extended Data Appendix B), electrical modulation in these experiments was delivered to a RNN after it had been trained, rather than during training when the network could learn to use it as a signal. In this way, our implementation of electrical modulation is more akin to experimental use of optogenetics in probing neuromodulation-dependent network behavior, or clinical administration of DBS and TMS in therapeutic modulation.

## Author contributions

B.T., S.C.P., K.M.T., H.T.S., and T.J.S. formulated the ideas; B.T., S.C.P. and T.J.S. designed the network architecture and simulation of the behavioral tasks; B.T. and S.C.P. implemented the design, ran the simulations, and carried out the data analysis; B.T., S.C.P, K.M.T., H.T.S., and T.J.S. wrote the manuscript.

## Acknowledgements

We thank Yusi Chen and Robert Kim for helpful discussions and Jorge Aldana for support with computing resources. Research was supported by the Kavli Institute for Brain and Mind at UC San Diego, ONR Grant N00014-16-1-2829, DARPA Grant HR0011-18-2-002, NIMH Grant R37-MH102441, and National Center for Complementary and Integrative Health Pioneer Award DP1-AT009925.

