## Supplemental data for "Neuromodulators generate multiple context-relevant behaviors in a recurrent neural network by shifting activity flows in hyperchannels"

### *Additional Materials and Methods*

#### Positive-Negative Output Task cont.

The positive stimulus consisted of an input of 0 from timesteps t=1 to t=50, an input of +1 from t=51 to t=75 (stimulus duration was 25 timesteps), and 0 for the remainder from t=76 to t=200. The null stimulus was an input of 0 throughout the whole trial (t=1 to t=200). The target outputs were as follows. For "positive output," the target output was 0 from t=1 to t=75 and +1 from t=76 to t=200. For "zero output," the target output was 0 throughout (t=1 to t=200). For "negative output," the target output was 0 from t=1 to t=75 and -1 from t=75 to t=200.

In the 9 behavior version all possible stimulus $\rightarrow$ output response pairs were added. As such, Behavior 1 was $+ \rightarrow$ negative output, $\emptyset \rightarrow$ negative output; Behavior 2 was $+ \rightarrow$ zero output, $\emptyset \rightarrow$ negative output; Behavior 3 was $+ \rightarrow$ positive output, $\emptyset \rightarrow$ negative output; ...; Behavior 9 was $+ \rightarrow$ positive output, $\emptyset \rightarrow$ positive output.

#### Comparison to context-dependent cued model cont.

To compare the neuromodulatory and context-cued RNNs tolerance to noise, we ran two types of simulation experiments. First, we added Gaussian noise of zero mean and standard deviation ranging from 0 to 4 to all input signals and measured RNN performance as the difference between target and actual output. At each level of noise, we simulated 4 independent models 100 times each and averaged their performances (Extended Data Fig. 1d). In a second experiment, after training, we simulated pure intrinsic network activity (no inputs). We again added Gaussian noise of zero mean and standard deviation ranging from 0 to 4. For each batch of 100 simulations on 4 replicate networks, we examined the consistency of final network states (t=200). To measure this, we computed the mean Euclidean distance of the 100 simulations final-time-step states from the centroid, giving a measure of final network state spread. For comparison of network types across different levels of noise, mean Euclidean distance was normalized to the no noise condition for the same network type. Larger mean Euclidean distance (higher spread) indicated more variable activity trajectories; lower mean distance (lower spread) indicated highly consistent activity trajectories (Extended Data Fig. 1e).

The phase portraits with flow fields were created by simulating a network to produce two state trajectories from distinct behavioral contexts. For this we used a neuromodulatory network ($f_{nm}$=9) and a context-cued model with persistent cues 0/+2.0. For each network configuration (with and without neuromodulator, and cued model), we computed the derivative of the neurons' rates across the trajectories to generate vector fields depicting the intrinsic flow fields of the network in the absence of any driving input. For the exogenously cued model, we then calculated the external drive required across the alternative cue-driven trajectory (cue=+2.0) to achieve the corresponding activity trajectory. We projected state trajectories and flow fields into the first three PCA space for visualization (Extended Data Fig. 1f).

We also assessed the effect of noise in the neuromodulator signal which we refer to as "intrinsic noise." We did this in two ways. First we trained replicate RNNs with added noise drawn from a Gaussian distribution with zero mean and set standard deviation. We then tested the RNNs across a range of intrinsic noise levels. We repeated this for RNNs trained with differing levels of intrinsic noise and again measured average performance (Extended Data Fig. 1g) and final state consistency (Extended Data Fig. 1h) exactly as was measured in Extended Data Fig. 1d,e. We also assessed variability in the neuromodulator signal a second way. Here we drew neuromodulator signals from a Gaussian distribution with a set standard deviation centered on a non-zero value (e.g. 2). We fixed these neuromodulator signals ("fixedstd" RNNs in Extended Data Fig. 1i,j);

each neuron was modulated by a different scalar (e.g. neuron 1 by 2.2; neuron 2 by 1.7, neuron 3 by 1.9, etc.). We then trained the RNN with this set of neuromodulator factors and then tested them across a range of random intrinsic noise levels drawn from Gaussian distributions. The same measures of performance and final state consistency are shown in Extended Data Fig. 1i,j.

##### Electrical modulation cont.

For graded electrical modulation, networks that did not achieve e-mod$_{50}$ at maximum stimulation (-9 units) were assigned a surrogate e-mod$_{50}$ value of -10 to calculate correlation to EC50. To account for possible missed correlation due to this substitution, correlation of EC50 to output at maximum modulation (-9) was also calculated (Extended Data Fig. 13c). For fully neuromodulated and electrically modulated networks (Extended Data Fig. 13d), we trained 30 RNNs with a factor of 9 and tested with four levels of electrical modulation (e-mod $\in \{-5.0, -1.0, 0.0, 1.0, 5.0\}$) in the absence and presence of neuromodulation.

#### Extended Data Appendix A

##### Relationship to isolated gain modulation

Gain modulation changes the slope of the unit activation function (the neuron's "intrinsic excitability") by changing a gain parameter $g$:

$$r = f(x; g)$$

where $r$ is the firing rate and $x$ is the synaptic current variable. For the sigmoid activation function this corresponds to:

$$r = \frac{1}{1 + e^{-gx}}$$

To see the effective change on the activation function of amplifying or dampening the weights we can compare the effect of gain modulation to weight modulation on the equation that governs neural dynamics:

$$\tau \dot{x}_i = -x_i + \sum_j W_{ji} r_{ji} + W_{ui} u + N(0, 0.1)$$

If we assume no input and no noise, we get a simplified equation describing neuron dynamics:

$$\tau \dot{x}_i = -x_i + \sum_j W_{ji} r_{ji}$$

Gain modulation $g$ gives:

$$\tau \dot{x}_i = -x_i + \sum_j W_{ji} r_{ji}(g)$$

while weight modulation with a neuromodulation factor $m$ gives

$$\tau \dot{x}_i = -x_i + \sum_j m \cdot W_{ji} r_{ji}(1)$$

To compare each type of modulation, we can consider the modified terms:

$$
\begin{aligned}
1 \cdot r(g) &= \frac{1}{1 + e^{-gx}} && \text{(gain effect)} \\
m \cdot r(1) &= \frac{1}{\frac{1}{m} + \frac{1}{m} \cdot e^{-x}} && \text{(weight effect)} \\
&= \frac{1}{\frac{1}{m} + e^{-x - ln\, m}}
\end{aligned}
$$

These effects are equivalent when

$$
g = -\frac{ln(\frac{1-m}{m} + e^{-x - ln\, m})}{x} \tag{1}
$$

So generally gain modulation, $g$, is only equivalent to a weight modulation by $m$ if the gain term is precisely the time varying function of $x$ defined by equation (1). Furthermore, for some values of $m$, there is no equivalent $g$ for certain values of $x$. E.g. for $m = 2$, $g$ is defined by equation (1) only for $x < 0$. For arbitrary values of $x$ with fixed, constant $g$ and $m$:

$$
r_{ji}(g) \neq m \cdot r_{ji}(1)
$$

except when $g = m = 1$, i.e., when there is no modulation. Thus, weight neuromodulation and gain modulation operate through different effects.

### Extended Data Appendix B

#### Chemical and electrical modulation

Chemical (neuromodulation) and electrical modulation in our network operate in directionally similar manner with qualitatively different effects. This can be seen by inspecting the equation governing each neuron i's synaptic current variable and thereby the network's activity dynamics:

$$
\tau \dot{x}_i = -x_i + \sum_j W_{ji} r_{ji} + W_{ui} u + N(0, 0.1)
$$

For any given neuron we can break up the terms by whether an input neuron is in the neuromodulated subpopulation or not:

$$
\tau \dot{x}_i = -x_i + \sum_k W_{ki} r_{ki} + \sum_q W_{qi} r_{qi} + W_{ui} u + N(0, 0.1)
$$

where $k$ is the index for non-neuromodulated neurons and $q$ is the index for neuromodulated neurons.

Neuromodulation in our model acts by scaling the target neurons outgoing weights by a factor $f$:

$$
\tau \dot{x}_i = -x_i + \sum_k W_{ki} r_{ki} + f \cdot \sum_q W_{qi} r_{qi} + W_{ui} u + N(0, 0.1)
$$

Electrical stimulation acts by adding exogenous synaptic current to target neurons. For a given neuron i in the non-neuromodulated subpopulation, electrical stimulation of the neuromodulated subpopulation is felt through altered incoming firing rates:

$$
\tau \dot{x}_i = -x_i + \sum_k W_{ki} r_{ki} + \sum_q W_{qi} r_{qi,Estim} + W_{ui} u + N(0, 0.1)
$$

and a neuron i in the neuromodulated subpopulation is additionally affected through direct stimulation:

$$\tau \dot{x}_i = -x_i + \sum_k W_{ki} r_{ki} + \sum_q W_{qi} r_{qi,Estim} + W_{ui} u + u_{Estim} + N(0, 0.1)$$

Thus we can see that both chemical and electrical stimulation act through the same term in the equation that governs neural synaptic currents yet in different manners. Neuromodulation directly scales presynaptic weighted inputs from neuromoduated neurons, whereas electrical stimulation acts by altering the firing rate of presynaptic neuromoduated neurons with an additional direct influence on the synaptic current if the neuron of interest is in the electrically stimulation subpopulation.

The similarities of these forms of modulation (acting through the same terms of the synaptic current equation) indicates why they can have similar affects on network output in some circumstances. Nevertheless, the differences in how they affect the synaptic current equation are propagated through the recurrent connections of the network at each time step which drives the distinct dynamical changes seen under chemical versus electrical modulation.

#### *Data and code availability*

The code and RNN models in this work will be made available at `https://github.com/tsudacode/neuromodRNN`

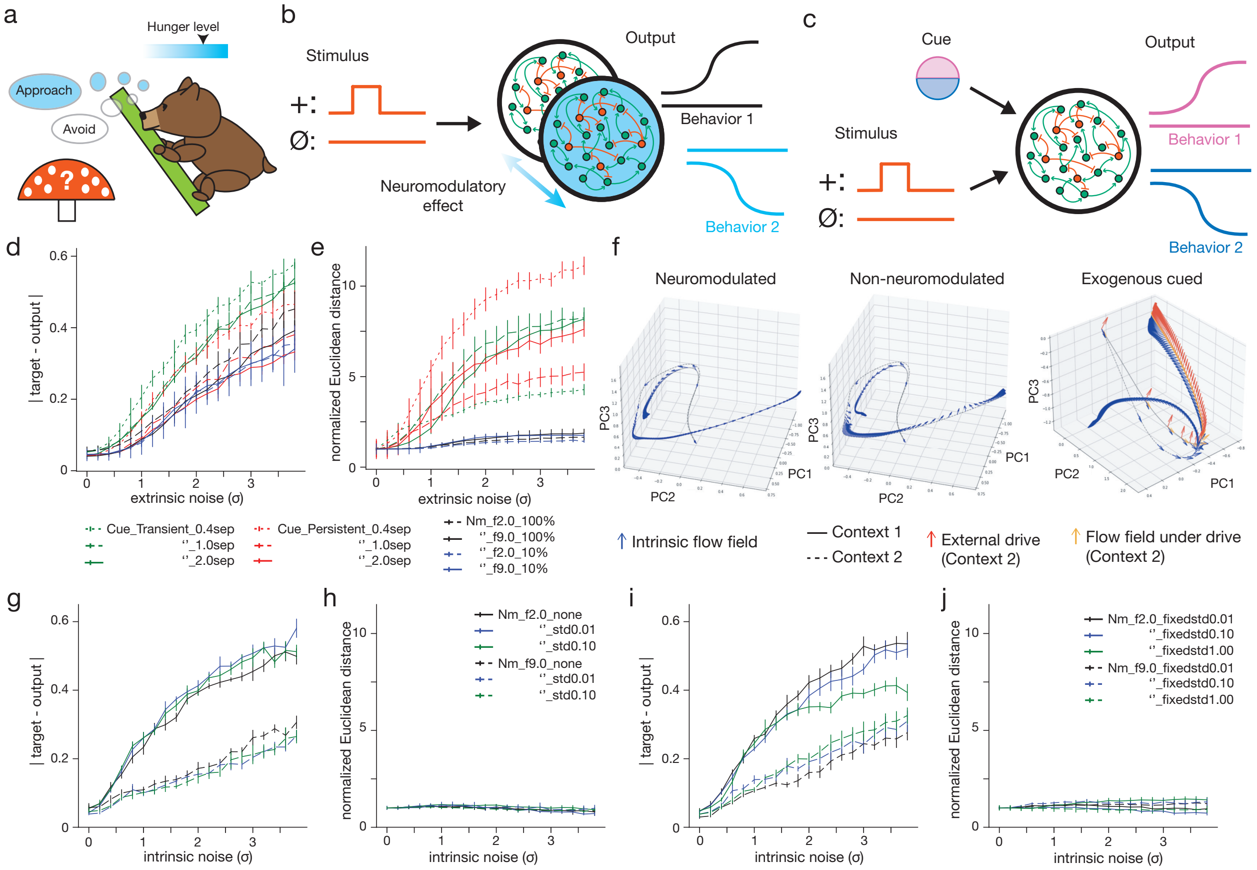


**Extended Data Fig. 1 | Noise robustness of neuromodulated and external cue-driven networks.** **a**, Neuromodulation is a fundamental biological mechanism, underlying e.g. how hunger level dictates a bear's behavior when encountering a mystery mushroom. **b**, Positive-Negative Output Task in neuromodulated network. Neuromodulatory effect alters network configuration to produce different behaviors. **c**, Comparison to alternative mechanisms of a context-cue input-dependent model. External cue is required to drive activity to generate different outputs. **d**, Output error for neuromodulated and exogenously cued models with increasing input noise. Noise is drawn from normal distribution with standard deviation (std) $\sigma$. Transiently cued models (green) — biologically more realistic than persistently cued (red) — have errors that increase more rapidly with increasing noise compared to neuromodulated (black—100% of network, blue—10% network; dashed lines factor 2.0, solid lines factor 9.0) or persistently cued models (red). **e**, Average Euclidean distance of neural trajectory endpoints between replicates when given no driving input across increasing levels of input noise. Neuromodulated networks (blue, black) generate network dynamics robust to noise, unlike cued models (green, red) which rapidly become unpredictable, exhibiting high variability — measured as increased Euclidean distance of trajectory endpoints on replicate trials — as noise levels increase. Cued models in **d**,**e** are highly dependent on specific cue values and separation amplitude between cues ("sep" in legend indicates input cue amplitudes separation). Figure caption continued on next page.

**Cont. Extended Data Fig. 1 | f**, Neuromodulation changes the flow field of the network in activity space (differences of vector fields in left and middle PCA activity plots). Blue vectors represent network's internal flow field (no driving input) along trajectories. Cue-based model (right) relies on external input (red arrows) to drive the network along desired trajectory in phase space (yellow arrows for Context 2 trajectory). For visualization clarity, flow field vectors indicate direction of activity change (without magnitude) and are shown only when network activity is changing more than a threshold of 0.02 units of Euclidean distance per timestep in full network activity space. **g**, Output error for models trained with intrinsic noise (i.e. noise in neuromodulator signal). Models trained with set level of noise (number after "std" in figure key indicates std of normal distribution from which noise was drawn) and then tested across spectrum of intrinsic noise ($\sigma$ is std of normal distribution from which noise was drawn). Of note, RNNs with higher neuromodulation factor were more robust in output error to increasing intrinsic noise, likely since relative noise-based deviation in neuromodulation factor from the mean was lower (compare a noise offset of 0.5 when the neuromodulation factor is 2 (25%) vs 9 ($\sim$5%)). **h**, Average Euclidean distance of neural trajectory endpoints between replicates with no driving input (same measure as **e**) across increasing levels of intrinsic noise. RNNs trained with intrinsic noise had neural trajectories robust to added intrinsic noise. Legend in **h** is for **g,h**. **i**, Same as **g** but for RNNs where neuromodulatory factors for each neuron in targeted population ($\boldsymbol{f_{nm}}$) are drawn from a normal distribution with mean and std indicated in legend ("fixedstd"). $\boldsymbol{f_{nm}}$ are fixed throughout training and testing. During testing intrinsic noise was added. Fixedstd RNNs show similar trend as those in **g**. **j**, Same as **h** but for fixedstd RNNs. Fixedstd RNNs also have neural trajectories robust to added intrinsic noise. Legend in **j** is for **i,j**. For **g–j**, error bars are SEM over 4 replicate models per condition.

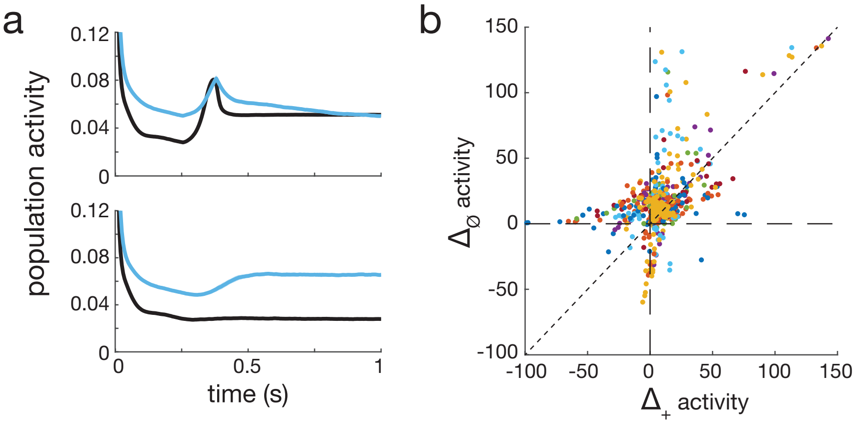


**Extended Data Fig. 2 | Neural activity under neuromodulation. a**, Mean whole network population activity for example RNN over 100 trials after training. Activity under neuromodulation (blue) is not simple transform of activity without neuromodulation (black). **b**, Difference of mean activity with and without neuromodulation ($\Delta$activity) on + vs $\emptyset$ stimulus trials for individual neurons. Each color represents an independently trained RNN (10 colors total). Points representing simple scaling of neural activity under neuromodulation would lie on dotted diagonal line.

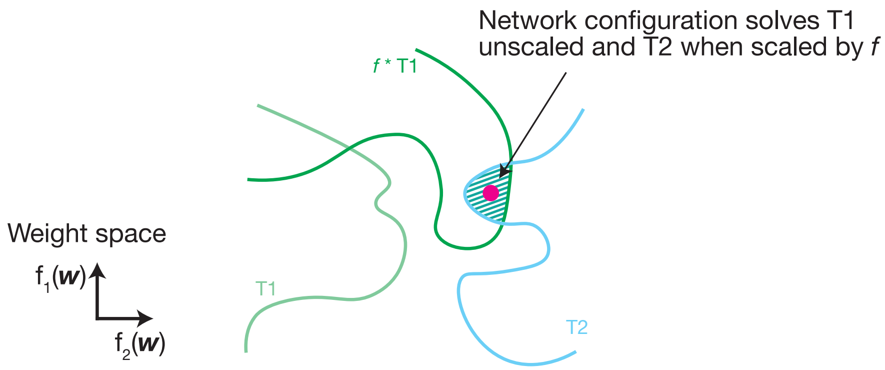


**Extended Data Fig. 3 | Relationship between neuromodulated weight configurations.** For the contradictory end behaviors in response to shared stimuli, as in the Positive-Negative Output Task, a single network without neuromodulation cannot simultaneously learn both tasks as depicted in this schematic by the lack of overlap between weight space that solves task 1 (T1) and task 2 (T2). By scaling weights in the network by a factor $f$, neuromodulation allows overlap between the $f \cdot$T1 and T2 spaces. The network solves task 1 when weights are unscaled (T1), and task 2 when weights are scaled ($f \cdot$T1 $\cap$ T2).

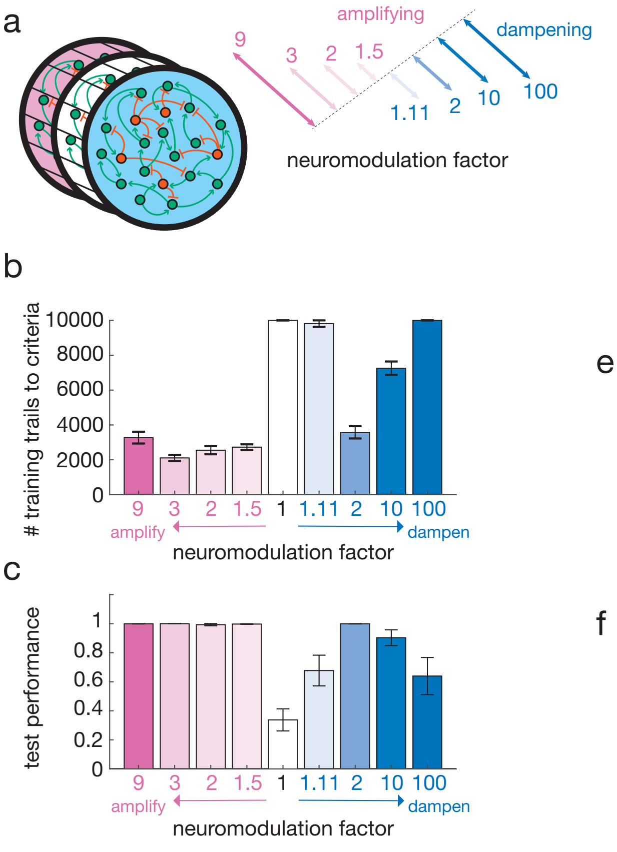


**Extended Data Fig. 4 | Neuromodulation weight scaling over a range of factors and weight types.** **a**, A range of neuromodulation factors were tested on the Positive-Negative Output Task. Factors were applied to weights initialized as described in Methods. **b**, All amplifying factors tested supported task learning in a similar number of training trials. Dampening factors that were either too small (e.g. 1.11) or too large (e.g. 100) led to longer training. **c**, All amplifying factors tested had perfect task performance. Dampening factors that were either too small or too large led to impaired performance, though better than without neuromodulation (factor=1). Extreme strong dampening effectively silences all transmission between neurons, impairing information flow in the network. Too little scaling (e.g. 1.11 factor dampening) did not create enough separation to distinguish the behaviors. **d**, Weight scaling modulation of input ($W_{in}$) and output weights ($W_{out}$) in addition to recurrent weights ($W_r$); of input weights $W_{in}$ only; and of output weights $W_{out}$ only. **e**, Like neuromodulation of $W_r$, neuromodulation of $W_{in}$, $W_r$, and $W_{out}$ at the same time led to successful training on the task after a comparable number of training trials. RNNs with neuromodulation of $W_{in}$ or $W_{out}$ alone were not able to fully learn the task before the 10,000 training trials stop criteria. **f**, Similar to neuromodulation of $W_r$ alone, neuromodulation of $W_{in}$, $W_r$, and $W_{out}$ at the same time led to near-perfect performance. RNNs with neuromodulation of $W_{in}$ or $W_{out}$ alone did not fully learn the task and performed at suboptimal levels. For **e** and **f**, all RNNs were trained with neuromodulation factor of 2. Error bars are SEM (10 independently trained RNNs for $W_r$, and 4 independently trained RNNs for other conditions).

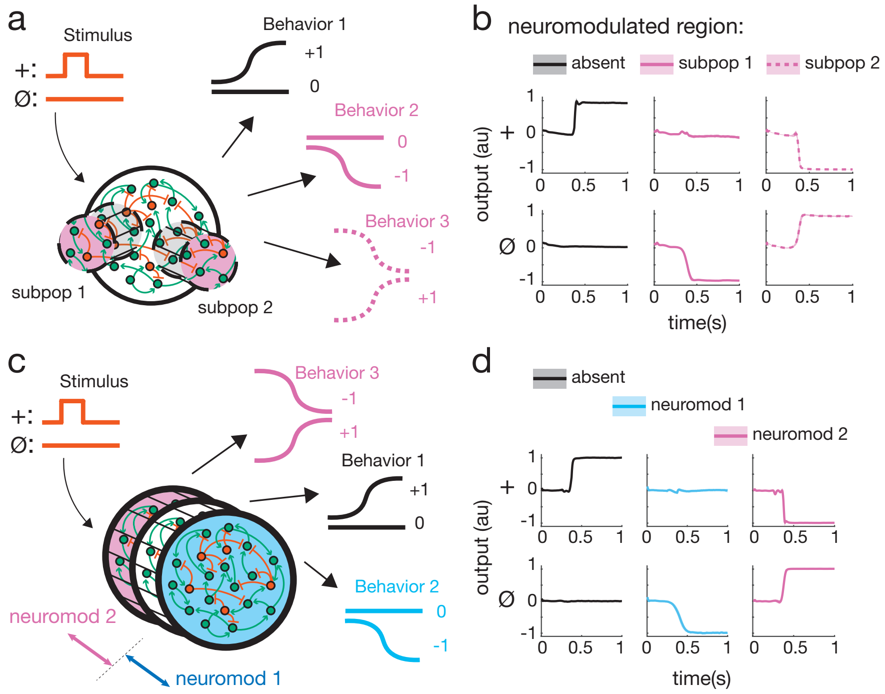


**Extended Data Fig. 5 | Multi-neuromodulator RNNs support multiple behaviors.** **a**, 3-behavior Positive-Negative Output Task. Neuromodulation (factor ($f_{nm}$)=2.5) of either subpopulations ("subpop") within a RNN (subpop1, subpop2) unlock unique behaviors (Behavior 2 (+ stimulus → 0 output, ∅ → -1), Behavior 3 (+ → -1, ∅ → +1), respectively). **b**, Multi-subpop targeted RNN from **a** successfully learns task. Top row: RNN output to + stimulus when different subpops are neuromodulated; bottom row: same for ∅ stimulus. **c**, Same task where different levels of neuromodulation (neuromod 1 ($f_{nm}$=0.5), neuromod 2 ($f_{nm}$=1.5)) are applied to whole RNN. **d**, Multi-neuromodulator RNN successfully learns task.

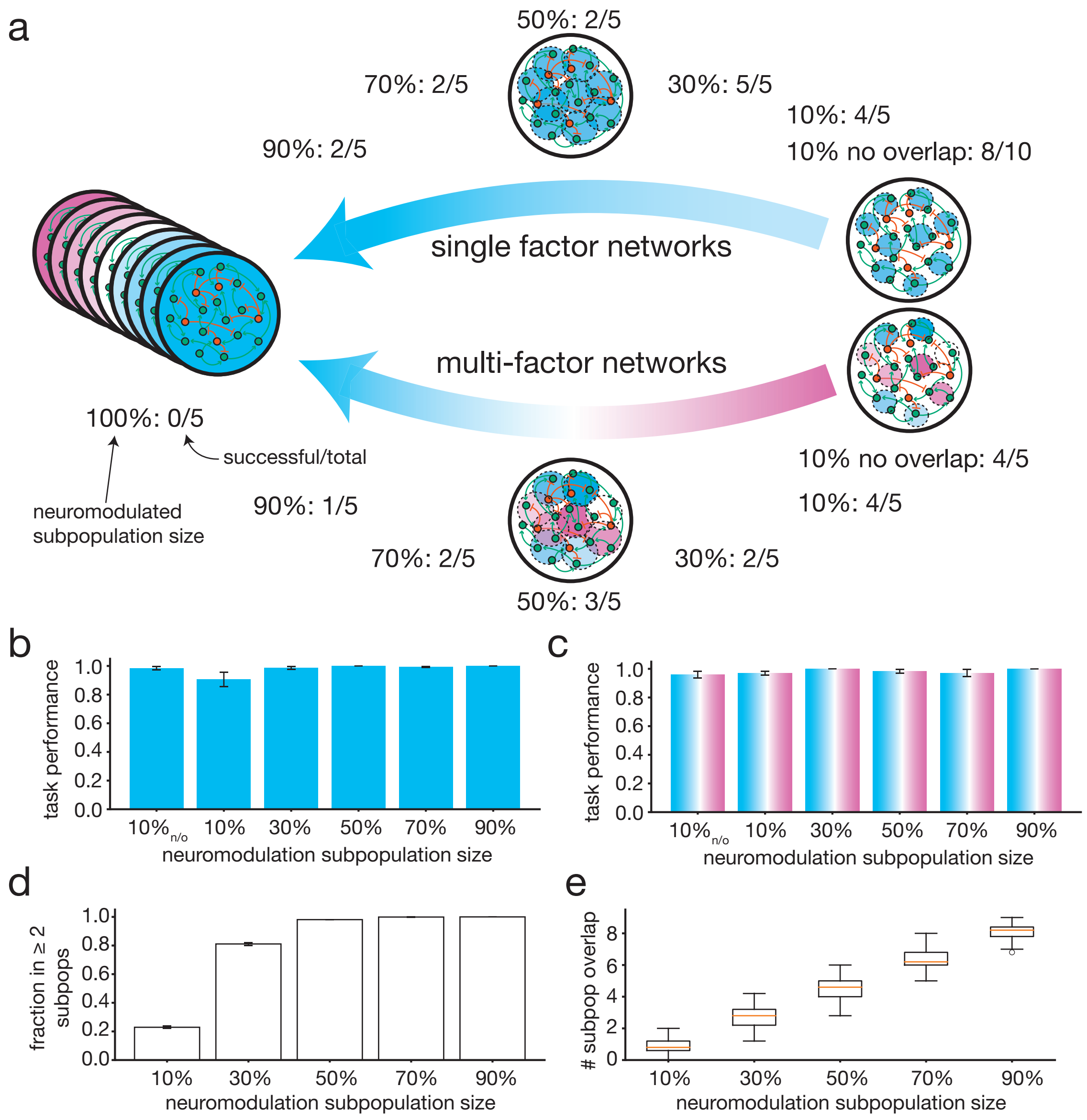


**Extended Data Fig. 6 | Spectrum of neuromodulation subpopulation size and factor. a**, Different conformations of neuromodulation of a single network support multi-behavior task (9-behavior Positive-Negative Output Task). Neuromodulation with a single factor (e.g. $f_{nm} = 2.5$) of non-overlapping and overlapping subpopulations across the spectrum of sizes could learn the full 9-behavior task, with larger overlapping subpopulations less consistently learning the full task ("successful/total" refers to independent networks that achieved successful training loss criteria over total attempted). Neuromodulation of the full network with different factors ($f_{nm} \in [1{:}1{:}9]$) consistently supported the 3 behavior task (5/5 successful/total), but not >3 behavior tasks (4-behavior 0/5; 5-behavior 0/5; 9-behavior 0/5 successful/total). Subpopulation neuromodulation with different factors ($f_{nm} \in [1{:}1{:}9]$) could also learn the 9-behavior task with overlapping and non-overlapping subpopulations, similarly exhibiting less consistent learning with larger overlapping subpopulations. **b**, Single factor networks test performance on 9-behavior task across networks that achieved training loss criteria. **c**, Same as **b** but for multi-factor networks. **d**, Fraction of neurons neuromodulated for ≥2 conditions across range of neuromodulated subpopulation sizes. **e**, Mean number of conditions a neuron was neuromodulated for across range of subpopulation sizes. For **d**,**e**, 5 replicates per condition. All error bars are SEM.

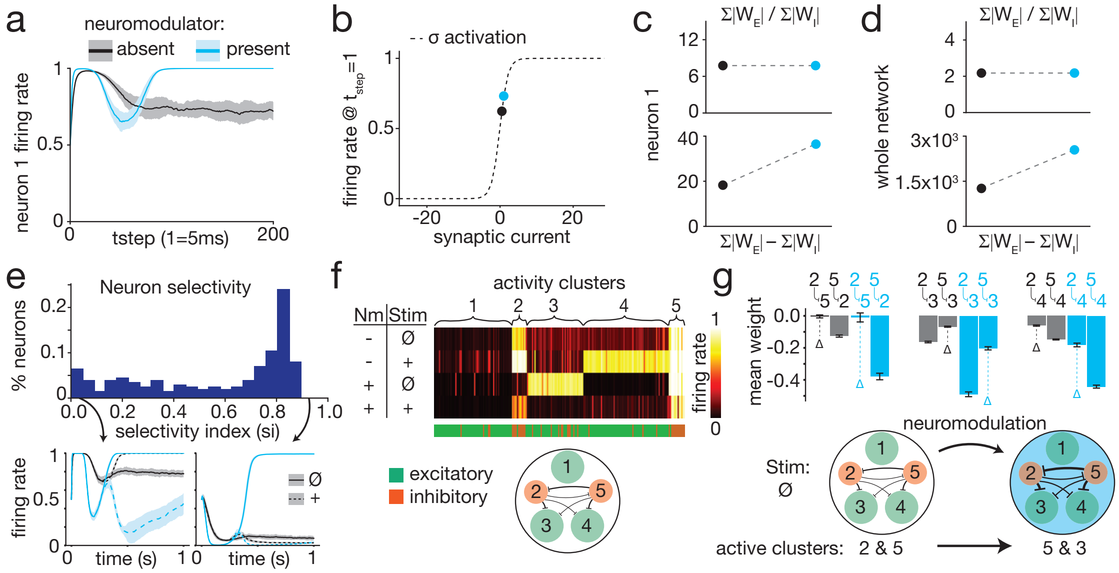


**Extended Data Fig. 7 | E-I difference scaling drives differential activity patterns.** **a**, Differential response of an individual neuron (neuron 1) in a whole-network neuromodulated RNN to same stimulus depending on neuromodulation presence. **b**, At the start of a trial, neuromodulation causes neuron 1 to receive different synaptic current input, shifting its firing rate. **c**, The different synaptic input under neuromodulation occurs due to amplification of the net difference in incoming excitatory and inhibitory weights; E/I balance is unchanged. **d**, E-I difference across the whole network is also amplified; E/I remains unaltered. **e**, Though all neurons in the RNN are influenced by the same neuromodulation, some exhibited activity selective for particular stimulus-neuromodulation combinations (high si; example neuron bottom right, si=0.89; see Methods); others were less selective (low si; example neuron bottom left, si=0.01). **f**, Neurons formed 5 clusters—2 predominantly inhibitory, 3 predominantly excitatory—whose activity coded for different stimulus-neuromodulation combinations. Each column of the heatmap is the mean firing rate of an individual neuron across conditions with excitatory/inhibitory identity labeled below. (Nm: neuromodulation present or absent; Stim: stimulus presented) **g**, Amplification of relative weight differences (Δs of mean weight) between inhibitory clusters drives cluster activity switches under neuromodulation: increased inhibition of clusters 2 and 4 by cluster 5 and disinhibition of cluster 3. In all panels light blue represents modulator present and grey/white modulator absent. Error bars are SEM.

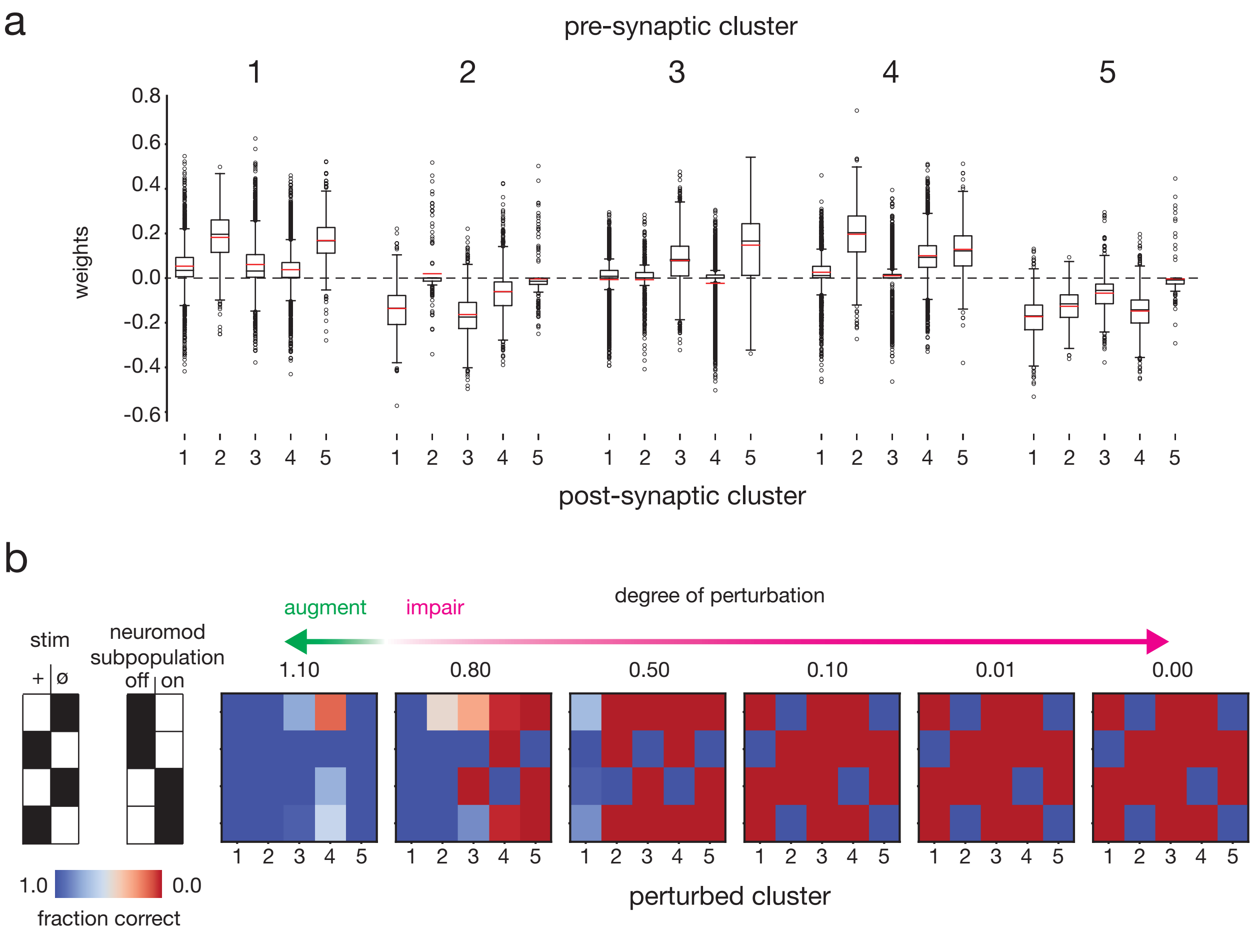


**Extended Data Fig. 8 | Inter-cluster weights and functional clustering for network in Extended Data Fig. 7. a**, Weight distribution from each cluster (pre-synaptic) to other clusters (post-synaptic). Clusters 2 and 5 were predominantly composed of inhibitory neurons resulting in inhibitory tone on downstream clusters, while clusters 1, 3, and 4 were predominantly excitatory. **b**, Augmentation (perturbation=1.1x) and impairment (i.e., synaptic lesions; perturbation $\in$ [0.80,0.50,0.10,0.01,0.00]x) of specific clusters led to impairment of performance. For most clusters, increased degree of synaptic impairment led to decreased performance, with higher sensitivity for stimulus-context associated clusters (Extended Data Fig. 7f; e.g. cluster 4 for + stim, neuromodulation off). In some stimulus-contexts, strong impairment or full lesion of clusters led to recovery of performance (perturbation $\leq$ 0.10 for clusters 2 and 5 with null stimulus without neuromodulation or positive stimulus with neuromodulation) suggestive of a counter-intuitive switch in underlying cluster dynamics.

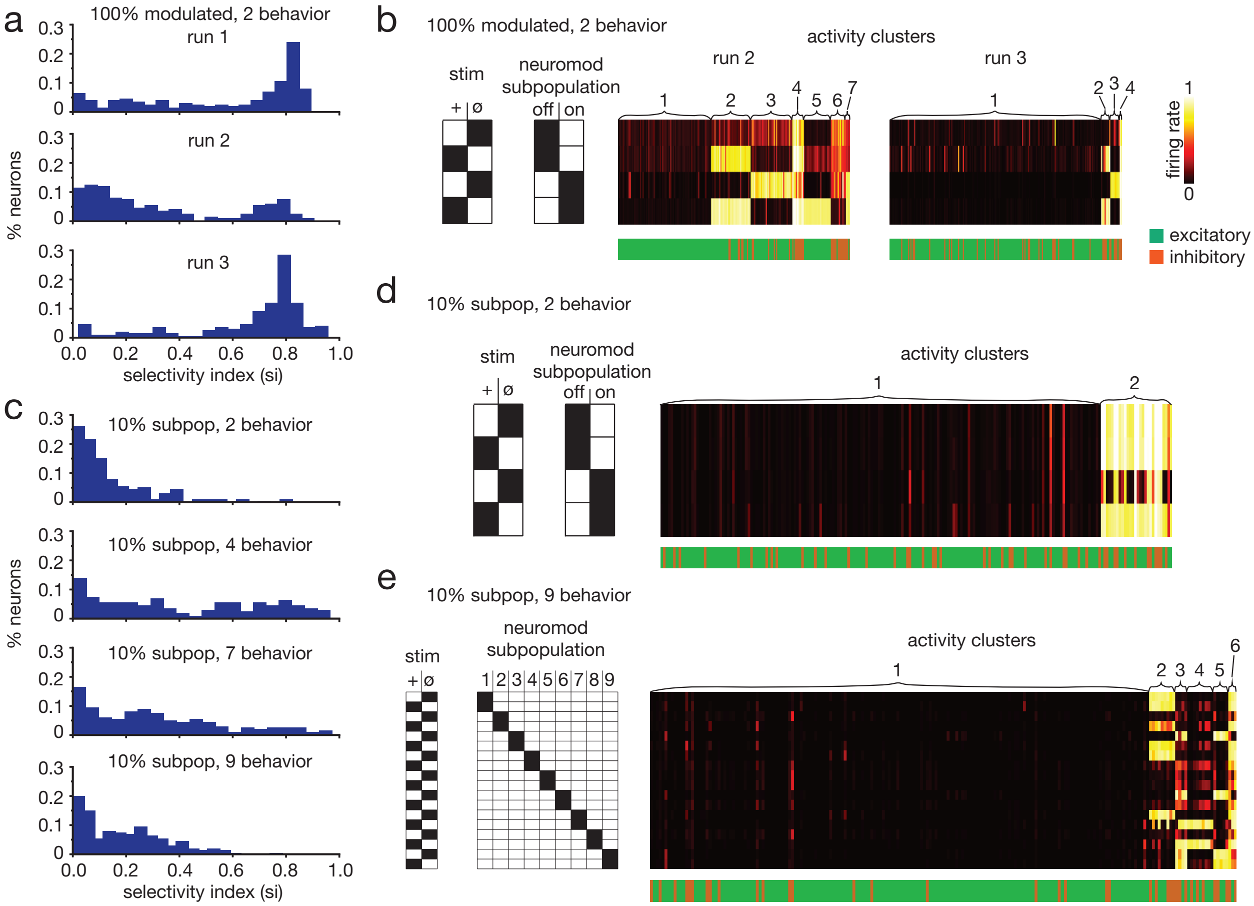


**Extended Data Fig. 9 | Independent networks have unique clustering patterns & different network configurations exhibit less selective neurons and complex, overlapping clustering profiles.** **a**, Neural selectivity profiles for 3 networks trained independently on the same condition (100% modulated, 2-behavior). All histograms exhibit high selectivity peak. **b**, Each network from **a** exhibits a unique neural activity clustering pattern including number of clusters and excitatory-inhibitory composition of clusters (see Extended Data Fig. 7f for run 1 cluster heatmap). **c**, Selectivity index histograms for example networks with different configurations and tasks. Compared to a network in which all neurons were neuromodulated trained on the 2-behavior Positive-Negative Output Task, network configurations with smaller neuromodulated subpopulations and more behaviors exhibited less selective neurons and more non-selective neurons. **d**, A network with 10% neuromodulated subpopulation trained on the 2-behavior Positive-Negative Output Task formed 2 activity clusters, showing high overlap across stimulus-context conditions. **e**, The stimulus-context clustering profile for a network with 10% neuromodulated subpopulations (non-overlapping) trained on the 9-behavior Positive-Negative Output Task. Cluster profiles are complex and highly overlapping. **d**,**e** cluster heatmaps and neuron type labels are same as **b**.

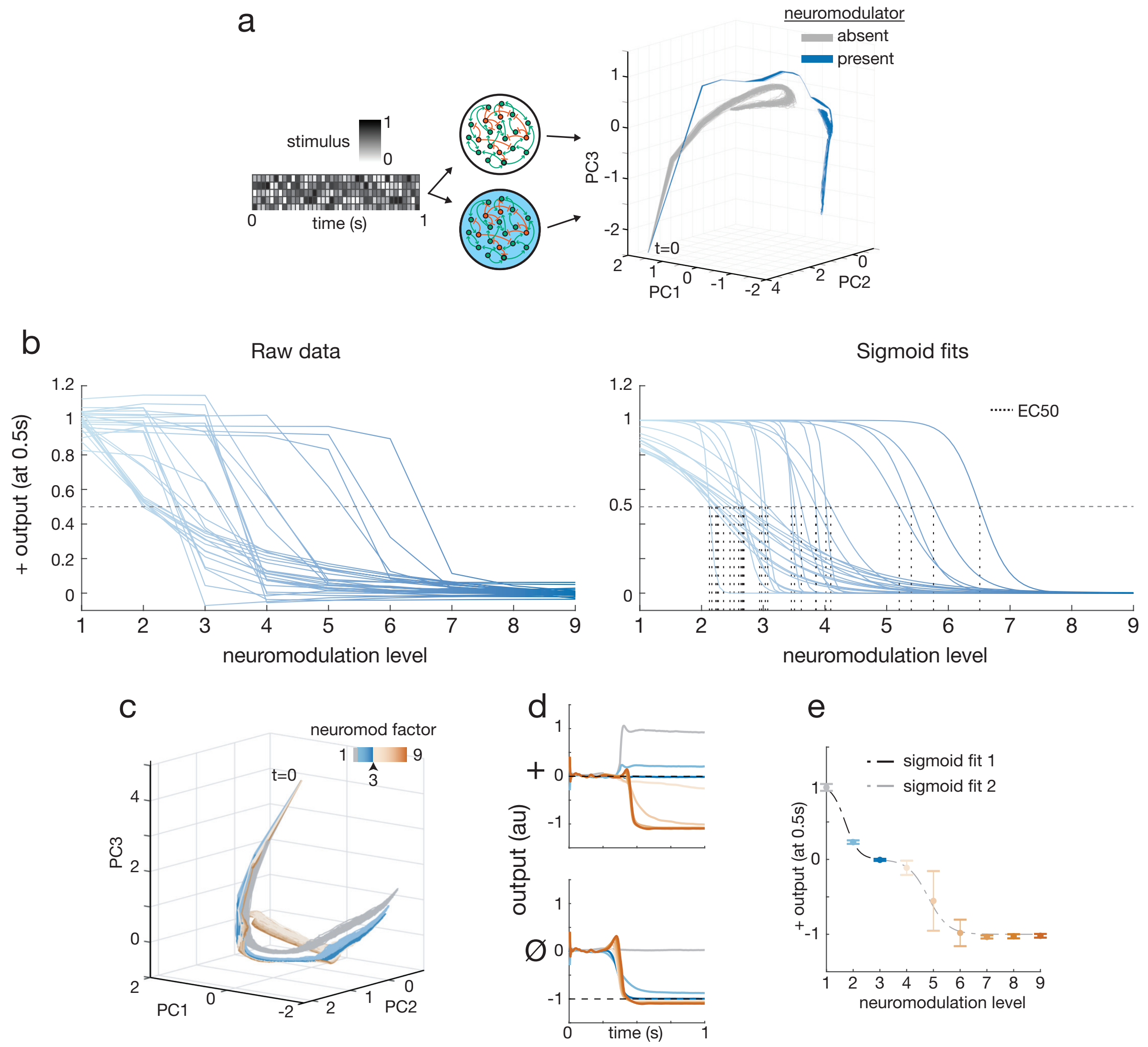


**Extended Data Fig. 10 | Variability in network neuromodulatory transitions.** **a**, Shotgun stimulus mapping of activity space inhabited by network in absence and presence of neuromodulation defines separable activity manifolds on which individual trajectories occur. **b**, Left: Output to positive stimulus at time point 0.5s of 29 networks at varying levels of neuromodulation. Each network was independently trained with amplifying neuromodulation factor 9 on the Positive-Negative Output Task and then tested at intermediate neuromdulation levels. Right: Sigmoid fits to raw data with EC50 of each curve indicated by dotted vertical line. EC50s ranged from 2.1 to 6.5. Slope of sigmoids ($\sigma$_slope) ranged from 0.9 to 26.3. **c**, For + stimulus, excessive neuromodulation pushes network activity into different space. **d**, Over-neuromodulation can drive inappropriate behavior (top) or no change (bottom). **e**, Transition dynamics for overmodulation is best fit by double sigmoid; first fits normal neuromodulation transition (blue) and second fits abberant neuromodulation transition (orange). For all PCA, top 3 PCs accounted for 80–92% of activity variance.

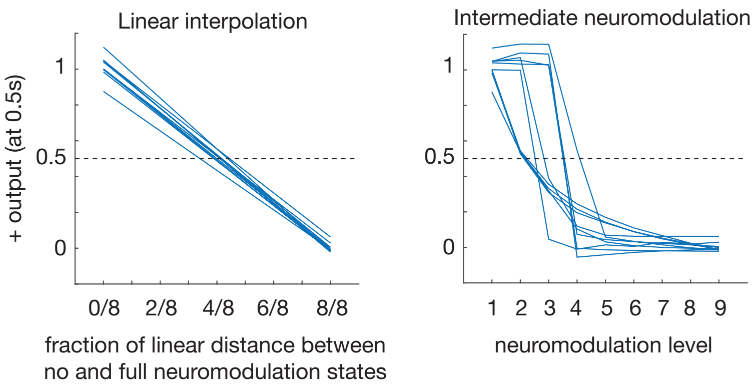


**Extended Data Fig. 11 | Unique geometry of transition manifold in phase space leads to highly variable network sensitivity (EC50).** Comparison of outputs for pure linear interpolation between no and full neuromodulation states (left) and intermediate neuromodulation levels (right) for same networks. Linear interpolation (left) gives linear output transition, with all networks tightly following a similar transition and intersecting half-maximal output (0.5) very close to the linear interpolation halfway point (4/8 distance). Intermediate neuromodulation (right) results in nonlinear output transition dynamic with high variability. For clarity, plots show only first 10 simulated networks (out of 29 total).

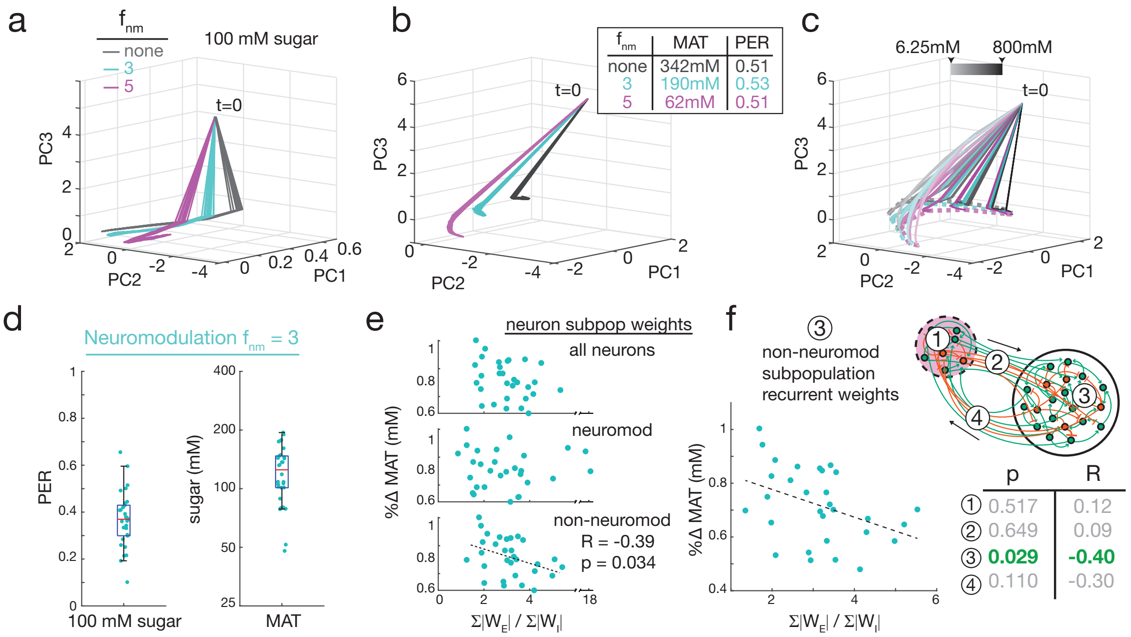


**Extended Data Fig. 12 | RNN models of *Drosophila* exhibit emergent transition behaviors and high variability of neuromodulator sensitivity.** **a**, For 100mM sugar, a RNN with intermediate neuromodulation ($f_{nm}$=3; cyan) generates activity between the neuromodulation extremes (none; grey and $f_{nm}$=5; magenta). **b**, Activity trajectory at MAT sugar concentration for intermediate neuromodulation of a RNN lies between those for neuromodulation extremes. **c**, Across sugar concentrations, intermediate neuromodulation trajectories (cyan) lay between neuromodulation extremes (grey and magenta), forming a 3-layer activity curtain ending on curved line (dotted lines) defined by trajectory endpoints across the sugar spectrum. **d**, Independently trained RNNs (n=30) exhibited high variability of PERs at 100mM sugar (0.10 to 0.66) and MATs (48 to 194mM). **e**, Normalized MAT change (%ΔMAT) vs E/I ratio for whole RNN and subpopulations. %ΔMAT had significant negative correlation to E/I ratio of the non-neuromodulated neurons ($p<0.05$, R=-0.39). **f**, %ΔMAT E/I ratio correlation was driven by recurrent weights within the non-neuromodulated subpopulation ③, rather than recurrent weights within the neuromodulated subpopulation ①, weights from neuromodulated to non-neuromodulated subpopulation ②, or weights from non-neuromodulated to neuromodulated subpopulation ④ as shown by Pearson correlation p-values (p) and coefficients (R) (for ①,②,④ correlation significance scores shown, scatter plots not shown).

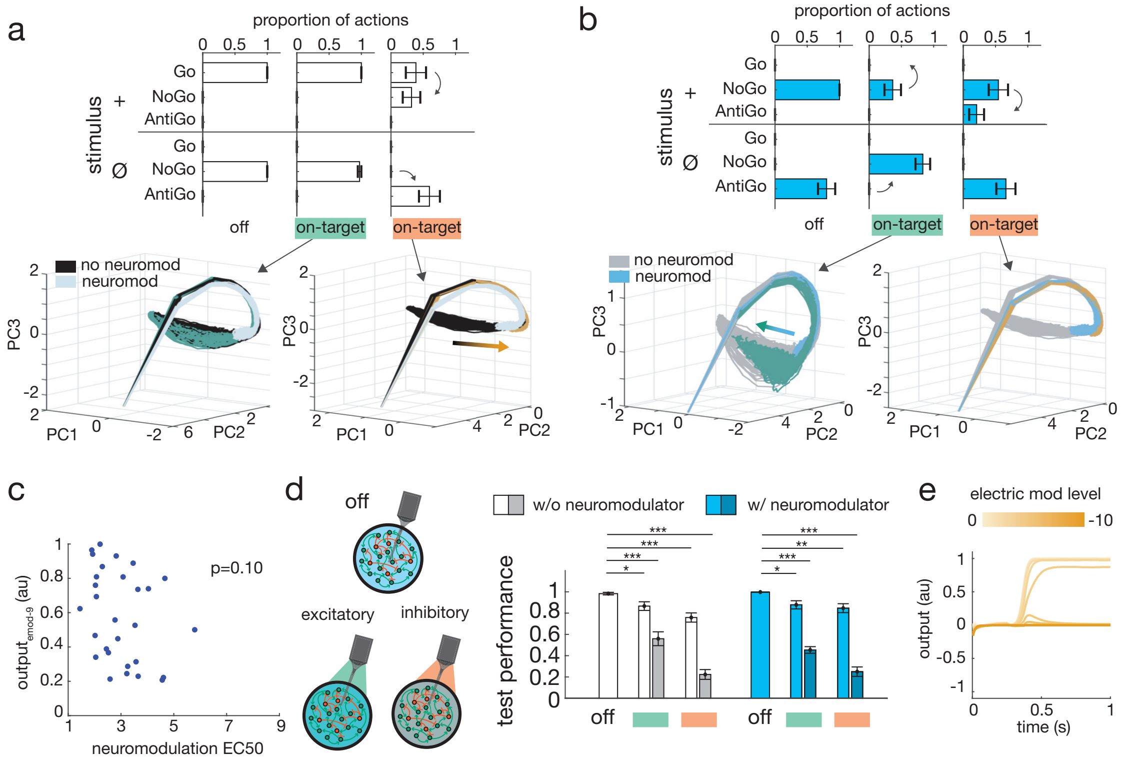


**Extended Data Fig. 13 | Targeted electrical modulation shifts network dynamics through independent circuit effect.** **a**, Behavioral and activity shifts in a RNN in absence of neuromodulation. Top: behavioral shifts with on-target e-mod. Bottom: activity dynamics under e-mod. Activity trajectories without e-mod are in black (neuromodulated trajectories in light blue for reference). E-mod can shift trajectories off black toward neuromodulated activity space (orange arrow) with corresponding shift of behavior toward neuromodulation-dependent behavioral set. **b**, Same as **a** with present neuromodulation. Top: behavioral shifts. Bottom: Activity trajectories without e-mod are in blue (non-neuromodulated trajectories in grey for reference). E-mod can shift trajectories from blue space toward non-neuromodulated activity space (green arrow) with corresponding shift of behavior toward non-neuromodulation behavioral set. **c**, Some networks did not reach an output level equivalent to half the response of maximal neuromodulation (EC50) even at the highest level of electrical modulation given of -9 units. To assess all networks electrical modulation sensitivity in comparison to neuromodulation sensitivity, network output at maximum electrical modulation is compared to EC50. Like EC50 vs e-mod$_{50}$, there is no significant correlation (p=0.10). **d**, E-mod of whole network in which whole network was targeted by neuromodulation ($f_{nm}$=9). Similar to RNNs with neuromodulated subpopulation with e-mod targeted to the same subpopulation, e-mod of fully neuromodulated RNN could impair performance. Bar graph shows two levels of e-mod (magnitude of input current 1 (lighter shade) or 5 (darker shade)) either excitatory (green indicator) or inhibitory (orange indicator) applied without (white/grey bars) or with simultaneous neuromodulation (blue bars). Error bars are SEM over 30 independently trained RNNs. *:p<0.01, **:p<0.001, ***:p<0.0001, Student's $t$-test. **e**, Example whole network-modulated RNN output in absence of neuromodulation to + stimulus with increasing inhibitory e-mod. Like subpopulation-targeted RNN in Fig. 5c, graded e-mod results in graded output transition, which varied from network to network.
